# Kite-shaped molecules block SARS-CoV-2 cell entry at a post-attachment step

**DOI:** 10.1101/2021.05.29.446272

**Authors:** Shiu-Wan Chan, Talha Shafi, Robert C. Ford

## Abstract

Anti-viral small molecules are currently lacking for treating coronavirus infection. The long development timescales for such drugs are a major problem, but could be shortened by repurposing existing drugs. We therefore screened a small library of FDA-approved compounds for potential severe acute respiratory syndrome coronavirus-2 (SARS-CoV-2) antivirals using a pseudovirus system that allows a sensitive read-out of infectivity. A group of structurally-related compounds, showing moderate inhibitory activity with IC_50_ values in the 1-5µM range, were identified. Further studies demonstrated that these ‘kite-shaped’ molecules were surprisingly specific for SARS-CoV and SARS-CoV-2 and that they acted early in the entry steps of the viral infectious cycle, but did not affect virus attachment to the cells. Moreover the compounds were able to prevent infection in both kidney- and lung-derived human cell lines. The structural homology of the hits allowed the production of a well-defined pharmacophore that was found to be highly accurate in predicting the anti-viral activity of the compounds in the screen. We discuss the prospects of repurposing these existing drugs for treating current and future coronavirus outbreaks.

## INTRODUCTION

The emergence of the coronavirus disease-2019 (COVID-19) has been a major global challenge that has led to unprecedented efforts to try to control the virus (1). These measures range from political and societal changes aimed at limiting virus spread, to attempts at eradication, as exemplified by vaccination. Whilst the former measures are highly unpopular, have serious impacts on economic factors and are of limited effectiveness, vaccination has so far proven to be highly efficient. Nevertheless, vaccine development and vaccination of global populations are lengthy processes, and new COVID variants may arise over the medium to long term that could require new vaccine development. This is particularly problematic with RNA viruses with high mutation rates, especially for vaccines that target the spike protein that is on the outside of the virion and thus subject to continuous selective pressure (2). For example the vaccine strain initially observed all over the world was replaced by the D614G spike variant in February, 2020 (3 months after the pandemic was announced) and other new variants are sweeping through the world currently (3–5) (6). Hence it seems appropriate to look for further control measures for the virus (7). Missing from the arsenal of effective measures to date has been an effective anti-viral therapy that could be administered prior to, or after acquiring the virus. So far, drug therapy has been limited to attempts to reduce the most life-threatening symptoms of the infection that arise due to over-stimulation of the immune response (8). We therefore lack a first line anti-viral defence to add to the current toolkit (9). Such first-line treatments would allow more time to develop new vaccines, could improve therapeutics and might be employed as prophylactics in those who cannot be vaccinated or do not respond well to vaccine. Such drugs, would ideally target conserved steps in the viral life cycle, be broad-spectrum and therefore generally applicable to COVID variants of the present and future (9).

The post-attachment entry step is one such conserved step (10). In order to enter host cells to initiate an infection, the virus must recognize and bind to a host cell receptor which then triggers virus-host cell membrane fusion to release the viral nucleocapsid into the host cell cytoplasm (11). Coronavirus spike protein is divided into an S1 attachment subunit and an S2 fusion subunit (12,13). The S1 subunits of the severe acute respiratory syndrome virus (SARS-CoV) and SARS-CoV-2 share 75% and 50% identity in the receptor binding domain (RBD) and the receptor binding motif whereas the more conserved S2 subunits share 88% and 100% identity in the fusion domain and fusion peptide (14). Receptor recognition is not conserved in coronaviruses. They use a range of host receptors. SARS-CoV-2 and SARS-CoV recognize the same receptor, the human angiotensin-converting enzyme 2 (ACE2), whereas the Middle East respiratory syndrome coronavirus (MERS-CoV) recognizes dipeptidyl peptidase 4 (DPP4) (12,13) (15) (16). The fusion mechanism, on the other hand, involving the formation of a 6-helix bundle, is conserved amongst viruses (10). Cleavage at S1/S2 and an internal S2’ site is a pre-requisite to prime fusion in coronaviruses (12,13,17). SARS-CoV-2 is unusual in that the S1/S2 boundary harbours a furin cleavage site so the spike protein is already cleaved in the mature virion (12). Viruses either fuse directly at the host plasma membrane under physiological pH or fuse at the endosome under acidic pH (10). Members of the coronavirus family can use either or both pathways (18). There is evidence to suggest that SARS-CoV-2 uses plasma membrane fusion as the default pathway but can use endosomal fusion if the plasma membrane protease, TMPRSS2, is not available; hence the micro-environment is important in dictating the entry pathway (19). However, it has been found that infection of ACE2-deficient lung cells depends on clathrin-mediated endocytosis and endosomal cathepsin L, indicating that endosomal fusion may well be the major entry pathway in a subset of cell types (20). Endosomal fusion is preceded by receptor-mediated endocytosis and trafficking to an acidic compartment to trigger fusion (10). Clathrin-, caveolae- and lipid-raft-mediated endocytosis have all been implicated in coronavirus infections (21). In SARS-CoV-2, both clathrin- and lipid-raft-mediated endocytosis have been demonstrated in two different 293T-ACE2 cell lines; despite somewhat contradictory results (22) (23). There is evidence that SARS-CoV-2 requires phosphatidylinositol 3-phosphate 5-kinase to traffick beyond the early endosome to reach the late endosome/lysosome for cathepsin L-catalysed S2’ cleavage to trigger endosomal fusion (17,23,24). Thus, the post-attachment entry steps depend heavily on a number of host signalling molecules which are amenable for drug targeting. Targeting conserved viral and/or host factors/processes negates the problematic drug escape mutants and is a current trend of generating broad-spectrum anti-virals (25).

Our aim is to find drug hits that target the entry steps, in particular the post-attachment step but any attachment blockers can be useful in virus-specific inhibition or universal synergistic inhibition with post-attachment inhibitors. Hence we employed a pseudovirus system in which the mouse leukaemia virus (MLV) is pseudotyped with the SARS-CoV-2 spike protein that would allow us to specifically screen for entry inhibitors (26) (27). During the current COVID crisis our first aim was to explore re-purposing of FDA-approved drugs and natural products, with the longer term goal of using any hits to generate a pharmacophore to inform next generation drug design.

## RESULTS

### 293T-ACE2 is a suitable cell type for pseudovirus drug screening

In order to re-purpose drugs for fast-tracking COVID-19 prophylaxis and treatments, we undertook screening of two libraries of FDA-approved drugs and natural products from APExBIO (28) by using MLV pseudotyped with the SARS-CoV-2 S protein (29), with the goal of targeting the major attachment and entry steps (Fig. 1). The spike protein is derived from the Wuhan-Hu-1 SARS-CoV-2 and has been codon optimized for mammalian expression (12). To find a suitable human cell type for the screening of drugs inhibiting viral infectivity we tested SARS-CoV-2-S, SARS-CoV-S, MERS-CoV-S and vesicular stomatitis virus (VSV)-glycoprotein (G) pseudoviruses against a range of cell types and employing the pseudovirus-encoded luciferase activity as a read-out for infectivity (Fig. 2). As expected, the control VSV-G pseudovirus, which has a broad host range (29), infected all cell types. SARS-CoV-S and SARS-CoV-2-S pseudoviruses did not infect the hepatocyte cell line, Huh-7. The heterogeneity in ACE2 expression in Huh-7 cell populations together with the widely varied characteristics of different laboratory-passaged Huh-7 lines may explain the discrepancy in the susceptibility of Huh-7 cells to native SARS-CoV-2 infection (13,30–32). In contrast, MERS-CoV-S pseudovirus, which preferentially binds DPP4 as a receptor rather than ACE2 (16), showed a high level of infectivity in Huh-7 cells. The green African monkey Vero cells (kidney), human colorectal epithelial Caco2 cells and human lung epithelial Calu3 cells all express a high level of ACE2 and are susceptible to native SARS-CoV-2 infection (31) but only Vero cells could be infected to a high degree by SARS-CoV-S and SARS-CoV-2-S pseudoviruses in this study. There was also a high background luciferase read-out from the empty (bald) pseudovirus in Calu3 cells. The human kidney epithelial 293T cells and human lung epithelial A549 cells express a low level of native ACE2 (31). A549 cells stably expressing recombinant human ACE2 showed SARS-CoV-S and SARS-CoV-2-S pseudovirus infectivity, but not to the same high level as 293T cells, which were therefore employed for initial drug screening. Compared to 293T cells not overexpressing ACE2, infectivity of SARS-CoV-2-S pseudovirus in 293T-ACE2 cells was about 100x higher (Fig. 3a,d). Quantitation by Western blot estimated the number of ACE2 receptors to be at least ten times higher in 293T-ACE2 cells than in the untransfected 293T cells with no loss of ACE2 expression in late passaged cells (P16) compared to early passaged cells (P4) (Fig. 3b,d). The number of trimeric spike proteins present on the surface of the pseudovirus in each infection experiment was also estimated (Fig. 3c,d). The data imply that the 293T-ACE2 cells’ ACE2 receptors will outnumber spike protein in the pseudovirus infection experiments (Fig. 3b,c,d), which is likely to be representative of the *in-vivo* situation, especially at early stages of infection. Furthermore the quantitation confirmed that the concentrations of proteins in the assays described below were well below the drug concentrations used. This was important to allow for the possibility of full inhibition by any given drug that was working by blocking the ACE2-Spike interaction and hence virus attachment to the cell.

**Figure 1:**
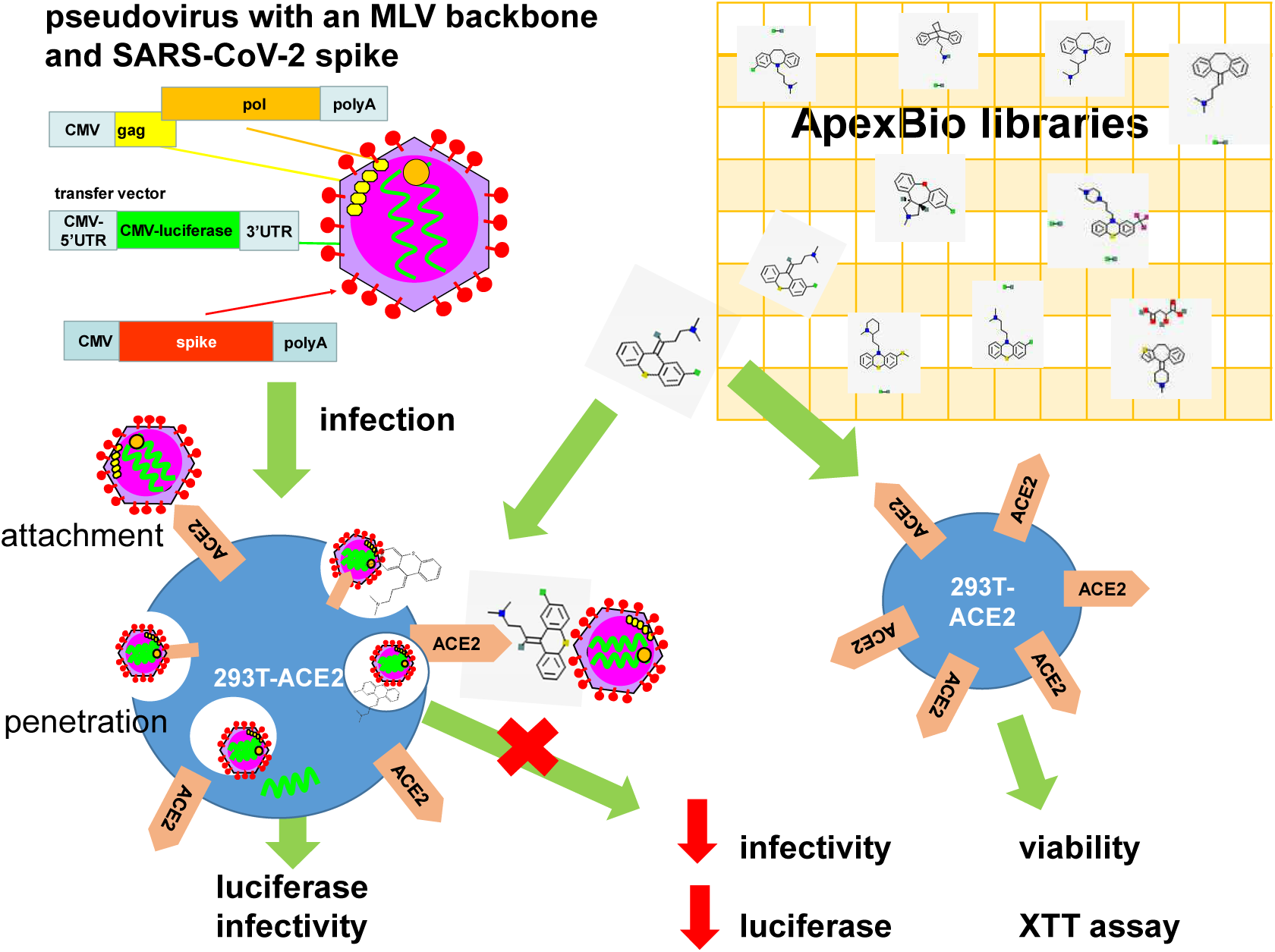
SARS-CoV-2 pseudotyped virus in anti-viral drug screening. Pseudovirus was generated on a mouse leukaemia virus (MLV) backbone using a three-plasmid system consisting of an expression vector for MLV *gag* and *pol*, a transfer vector carrying a luciferase reporter gene and an expression vector encoding the SARS-CoV-2 spike protein. Pseudovirus was used to infect 293T cells stably expressing the human angiotensin-converting enzyme 2 (ACE2). Pseudovirus entry was mediated by the binding of the spike protein to the ACE2 which then undergoes receptor-mediated endocytosis to trigger endosomal fusion to release the luciferase reporter gene into cell cytoplasm. Infectivity was measured as luciferase read-out. Drugs from two APExBIO libraries were screened for their ability to inhibit infectivity by measuring the reduction in luciferase activity. Since the spike protein only mediates virus entry, the pseudovirus system could be used to identify drug hits that inhibit SARS-CoV-2 entry steps only. Drug cytotoxicity was measured using an XTT viability assay in non-infected cells. Drug images were obtained from PubChem and APExBIO.

**Figure 2:**
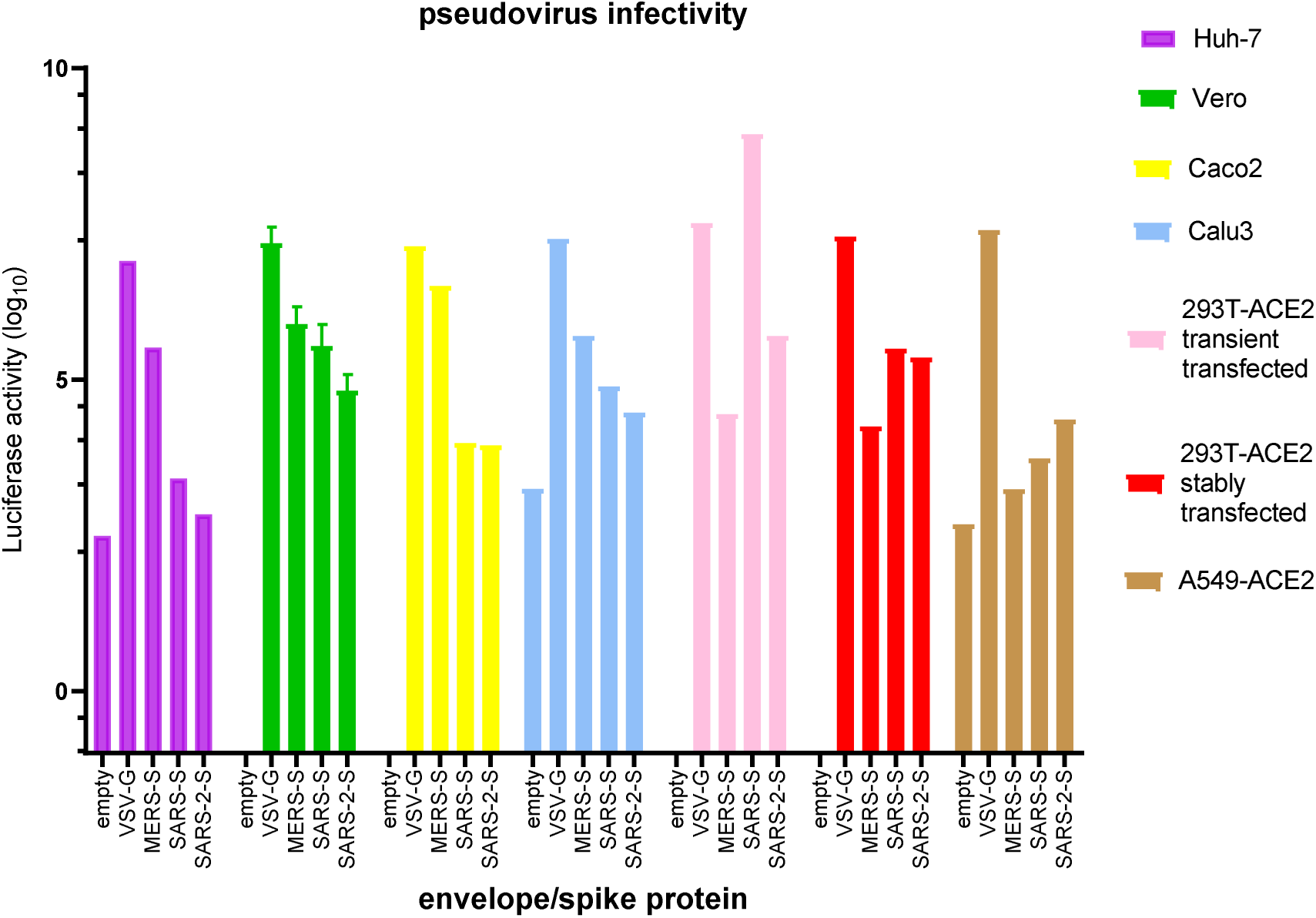
Infectivity of pseudoviruses in a range of cell types. Mouse leukaemia virus pseudotyped with no envelope protein (empty), glycoprotein from vesicular stomatitis virus (VSV-G) and spike protein (S) from Middle East respiratory syndrome coronavirus (MERS-S), severe acute respiratory syndrome coronavirus (SARS-S) and SARS-2-S was used to infect a range of cell types, as indicated, in 24-well plates for 72h. Infectivity was measured as luciferase activity. Data represent the mean of two repeats for Vero cells and one experimental result for the other cell types.

**Figure 3:**
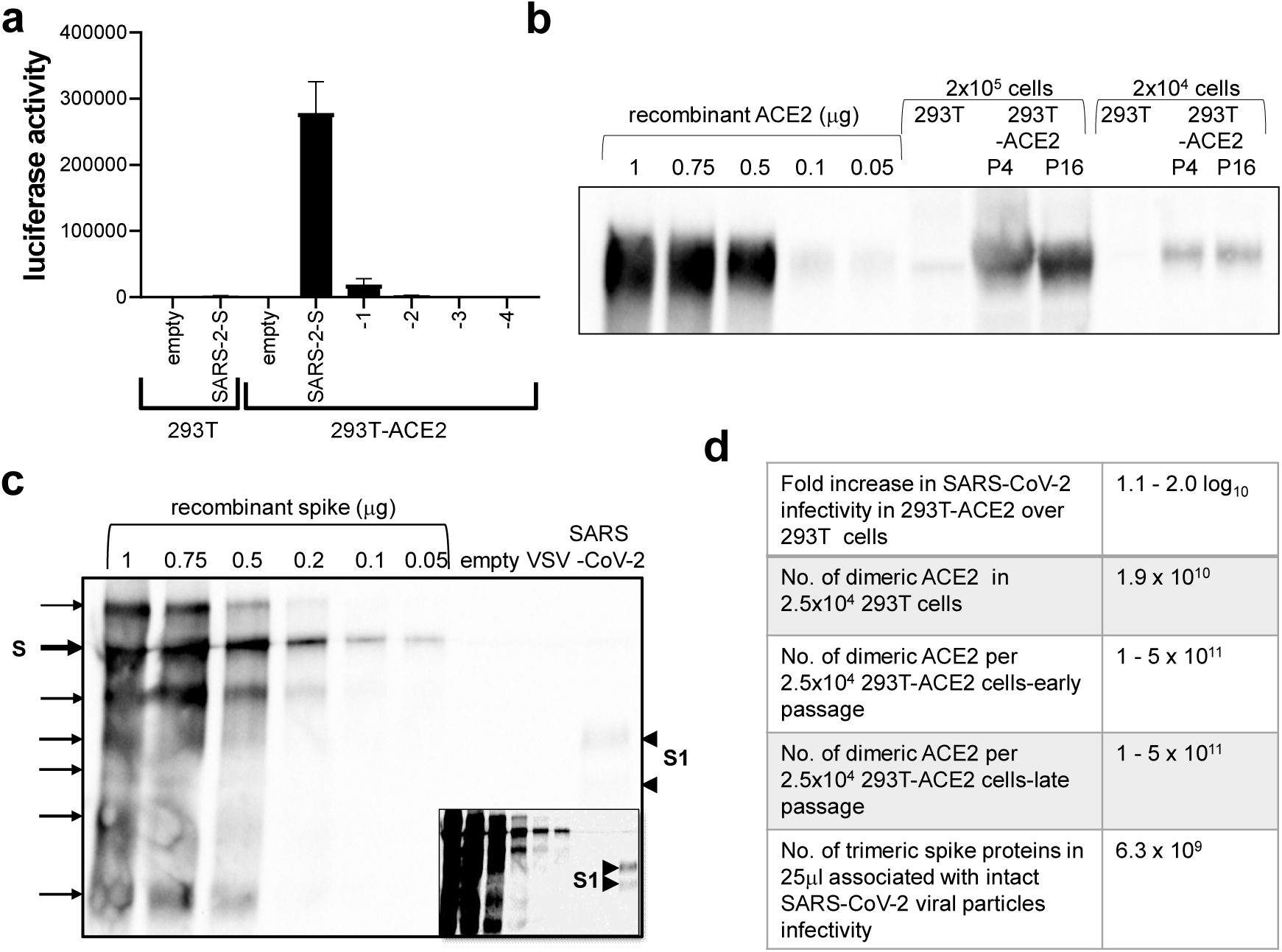
Quantification of ACE2 and spike protein. (a) Serial ten-fold dilutions of the SARS-CoV-2-S pseudotyped virus were used to infect 293T-ACE2 cells. Luciferase activity was measured at 72h and compared to that of 293T cells infected with undiluted pseudovirus. (b) Western blot of ACE2 from 2.5×10^6^ and 2.5×10^4^ 293T cells and 293T-ACE2 cells from early (P4) and late (P16) passages. The protein bands were quantified against a standard curve of recombinant ACE2. (c) Western blot of spike protein from 10μl of empty, VSV-G and SARS-CoV-2-S pseudovirus particles. The protein bands were quantified against a standard curve of recombinant spike protein. The spike protein in the SARS-CoV-2-S pseudovirus has been cleaved to yield the S1 subunit. The inset shows the same blot at higher contrast for clarity. Low exposure blot was used in quantification. The recombinant protein is near full-length and has been stabilized with the removal of the furin-cleavage site and exhibits many glycosylated and degraded forms. (d) A table summarizing the calculations. After estimating the µg of ACE2/spike proteins from the standard curve, the number of molecules was calculated by converting µg into moles multiplied by Avogadro’s number. The range reflects data calculated from 2.5×10^6^ and 2.5×10^4^ cell loading. The number of spike proteins was adjusted using the assumption that 76% of the spike protein are not associated with viral particles (they are secreted or degraded virion associated with extracellular vesicles). Amongst the viral particles, 80% are empty viral particles (non-infectious, no genome but retain spike) and the rest of the 20% intact particles had 0.4% infectivity (85).

### Kite-shaped molecules inhibit SARS-CoV-2 pseudovirus infection

An initial screen of a library of 1363 FDA-approved drugs was carried out using the above-mentioned cell line. The drugs at 10μM were incubated with the cells after dilution of the drugs into cell growth media from (predominantly) DMSO-solubilised stock solutions or (occasionally) ethanol-based or water-based stocks. Any cytotoxicity effects of the drugs at this concentration were controlled for using an XTT cell viability assay. After screening one-third of the compounds it became apparent that there was a prevalence of inhibitory activity found within a class of molecules that displayed a similar structure and a shape reminiscent of a traditional Chinese Kite. These had a well-conserved tri-cyclic core structure (forming the sail of the kite) and a more variable extension from the central 6- or 7-membered ring (forming the tail of the kite). We, therefore, selected 61 kite-shaped molecules from the two libraries. Five that were cytotoxic were excluded at this stage; the remaining molecules showed a range of activity against pseudovrius infectivity. Supplementary Table S1 summarises the experimental data for the kite-shaped molecules.

### Kite-shaped molecules specifically inhibit SARS-CoV-2 pseudovirus infectivity

Because the kite-shaped molecules could potentially inhibit both SARS-CoV-2-specific entry steps and MLV-mediated post-entry steps or the reporter, we tested the top eight hits against MLV pseudotyped with SARS-CoV-2-S, SARS-CoV-S, MERS-CoV-S and VSV-G, which share common MLV post-entry steps but differ in receptor recognition and entry mechanisms (18,33). We also included the 14^th^ ranked hit, trimipramine, because it had previously been identified as a specific SARS-CoV entry blocker and an inhibitor of SARS-CoV-2 infection (26). All the selected kite-shaped molecules showed effects similar to those of hydroxychloroquine, a known blocker of SARS-CoV-2 entry, by specifically inhibiting infectivity of the SARS-CoV-2-S, SARS-CoV-S, MERS-CoV-S but not VSV-G pseudotyped viruses, suggesting that the nine kite-shaped molecules target entry steps specific to these three coronaviruses (Fig. 4a). The kite-shaped molecules inhibited SARS-CoV-S and SARS-CoV-2-S pseudoviruses equally well and the inhibition was 1.2 to 8-fold higher than that for MERS-CoV-S pseudovirus, suggesting that they may, in addition, discriminate between different receptor-mediated pathways for viral entry. Although low levels of inhibition of VSV-G pseudovirus infectivity were indicated for a few of the nine compounds, this may be due to some cytotoxicity at 10μM, as suggested by the correlation between % inhibition and % viability (Fig. 4b). Alternatively, it could be due to inhibition of common post-attachment pathways in late endosome/lysosome. Hydroxychloroquine, a lysosomotropic agent, did not inhibit VSV-G pseudovirus infection, where fusion takes place at the early endosome stage (17). In contrast, the reverse transcriptase inhibitor, tenofovir disoproxil fumarate, completely inhibited infection of all pseudoviruses. These results imply that the kite-shaped molecules target the SARS-CoV-2 entry steps specifically, rather than any post-entry steps mediated by the MLV or the reporter.

**Figure 4:**
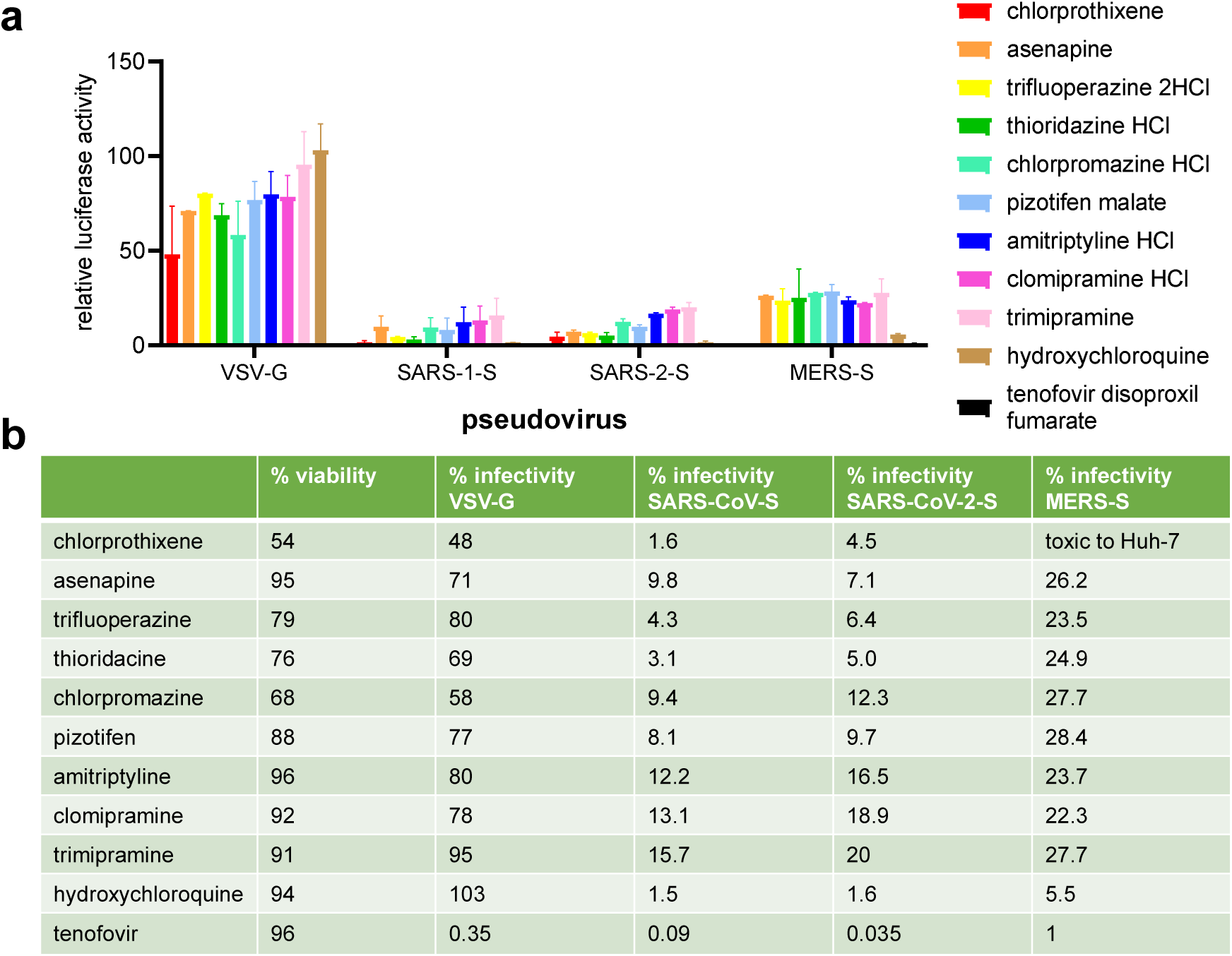
Kite-shaped molecules specifically inhibit coronavirus infection. Mouse leukaemia virus pseudotyped with glycoprotein from vesicular stomatitis virus (VSV-G) and spike protein (S) from severe acute respiratory syndrome coronavirus (SARS-S), SARS-CoV-2 (SARS-2-S) and Middle East respiratory syndrome coronavirus (MERS-S), was used to infect 293T-ACE2 cells, in a 96-well plate for 48h in the presence of the drug, as indicated, with 1h pre-treatment. (a) Infectivity was measured as luciferase activity and expressed as % infectivity versus infected, DMSO solvent control. Date are presented as mean +/− SD of two repeats. (b) Mean % infectivity was tabulated together with % viability.

Although the A549 cells transfected with ACE2 showed less propensity for infectivity than the HEK293T cell line (hence noisier luciferase readouts), a test of eight of the above-mentioned compounds with the A549-ACE2 system also demonstrated good inhibition of SARS-CoV-2 spike-mediated infectivity (Supplementary Information Fig.S1). Of the tested kite-shaped compounds, only asenapine failed to prevent infectivity in the A549-ACE2/pseudovirus system; whilst chlorprothixene, thioridazine and pizotifen malate showed the strongest inhibitory activity. As expected, tenofovir showed complete inhibition of infectivity in this lung-derived cell line, although like asenapine, hydroxychloroquine appeared to lack inhibitory action. Overall these results suggest that both airway and kidney cells expressing ACE2 can be treated with kite-shaped inhibitors with the proviso that cell-specificity may be a factor for some of the compounds tested.

### Efficacy of the kite-shaped molecules

To deduce the efficacy of the kite-shaped molecules, we generated dose-response curves using a range of drug concentrations from 10μM to 0.05μM and using hydroxychloroquine for comparison (Fig. 5a). The kite-shaped molecules displayed IC_50_ values from 1.9μM to 4.7μM, compared to 0.7μM for hydroxychloroquine (Fig. 5b). All the drugs showed *de minimis* cytotoxicity at 5μM and even at the highest drug concentration employed, the cells generally displayed a viability above 76% (chlorprothixene and chlorpromazine are exceptions with cell viability at 54% and 68%, respectively (Fig. 4b, Fig. 5). Since we diluted the drugs in cell growth medium, a confounding effect on the determination of IC_50_ could be water solubility of the drugs. Most of the drugs are readily water soluble in their charged state (pizotifen malate is the exception), but they may partition into the cell membrane via their uncharged forms which will have very low water solubility (Supplementary Table S2).

**Figure 5:**
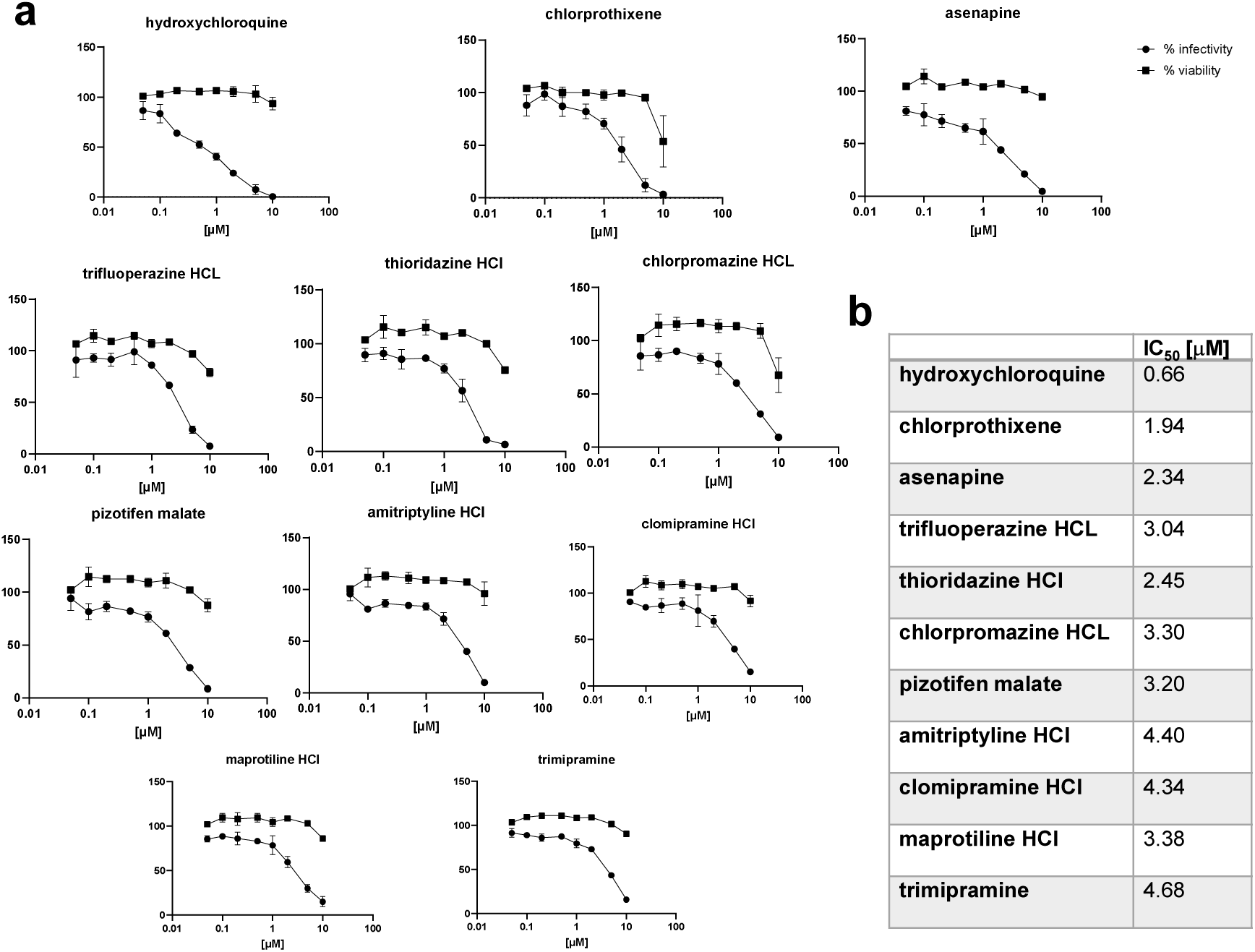
Dose-response curves of kite-shaped molecules in SARS-CoV-2-S inhibition. Mouse leukaemia virus pseudotyped with spike protein (S) from severe acute respiratory syndrome coronavirus-2 was used to infect 293T-ACE2 cells in 96-well plates for 48h in the presence of serial doses of the drug, as indicated, with 1h pre-treatment. (a) Infectivity was measured as luciferase activity and expressed as % infectivity to infected, own solvent control (dimethylsulphoxide, ethanol or water). Viability was measured by XTT assays in un-infected cells and expressed as % viability to solvent control (dimethylsulphoxide, ethanol or water). Data are presented as mean +/− SD of two repeats. (b) Summary of IC_50_ values.

The similarity in the overall structure of the kite-shaped molecules allowed the generation of a pharmacophore (Figure 6). Pharmacophores for tricyclic antidepressants (TCAs) have previously been described, showing the importance of the two outer aromatic rings, one of which is more hydrophobic. The tail region in the pharmacophore shows H-bonding propensity and the ability to form a positive charge on an amine group (34) (35). The pharmacophore generated from the SARS-CoV-2 infectivity assay displayed similar features to these prior studies (Figure 6a) but with more tightly defined distances and angles between the three main pharmacophore features. The three-feature minimal model was able to classify active compounds with predictivity values above 90%. When the complete continuous data was split into active and non-active compounds based on log activity values, the model still showed F score values of 70%. The model showed excellent predictive ability with both training and complete datasets (Figure 6b,c).

**Figure 6:**
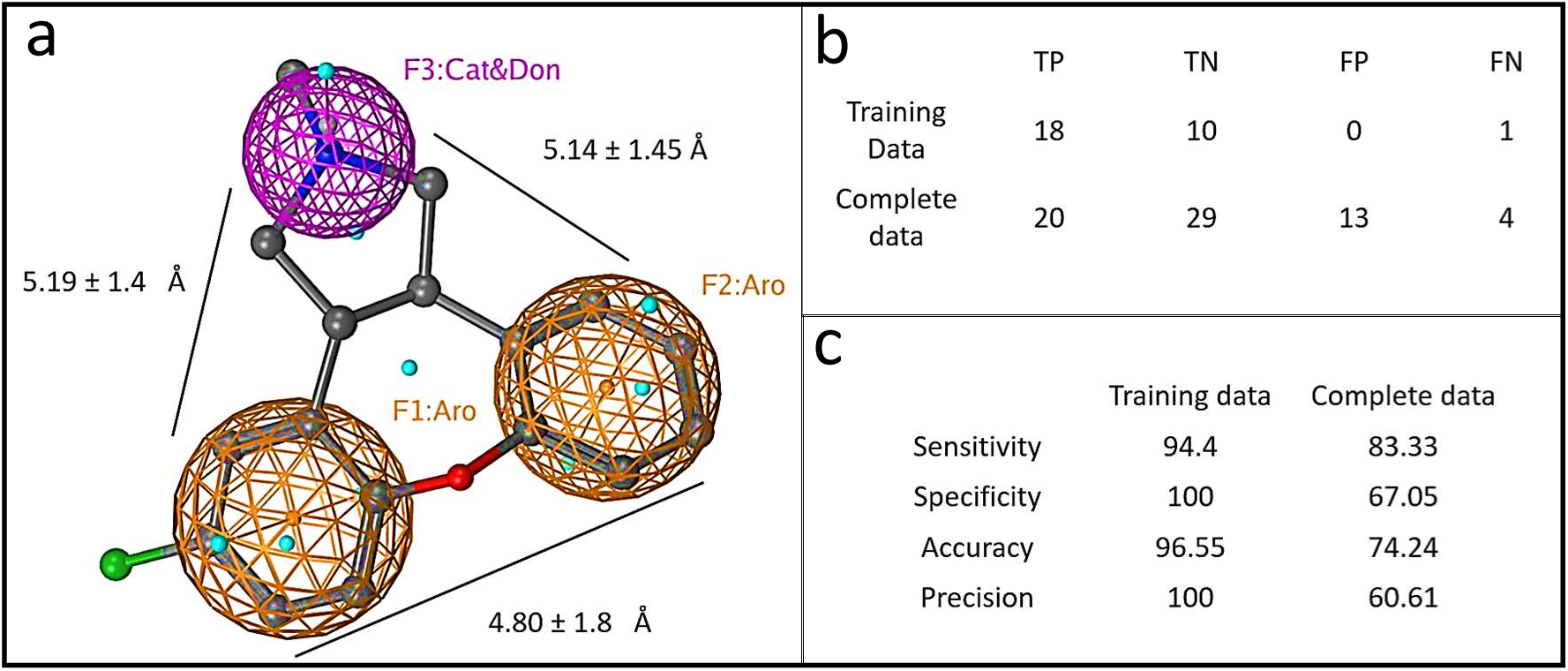
Graphical depiction of pharmacophore model. a) A three point pharmacophore model based on the kite-shaped molecules. The asenapine structure is superimposed (ball and stick representation) for comparison. Brown mesh represents aromatic moieties (Aro) and magenta mesh represents a H-bond donor/cation group (Cat&Don). Small spheres (cyan) highlight features in asenapine that are not relevant for the overall pharmacophore. (b) Displays the numbers of true (T) and false (F) positive (P) and negative (N) hits within the datasets that are discriminated by the pharmacophore (see Methods). Panel (c) summarises the pharmacophore model performance.

### Kite-shaped molecules inhibit pseudovirus entry

To study the mechanisms of inhibition in greater detail, we undertook a time-of-addition experiment in which drugs were added at different time-points during infection (Fig. 7a) with the hypothesis that the time-point(s) may distinguish different step(s) that may be inhibited by the drugs. As before, we studied the top nine hits together with trimipramine. The kite-shaped molecules were able to inhibit infection when added during the entry step (with and without 1h pre-incubation), to similar extents to the full-treatment (Fig. 7b). Most of the kite-shaped molecules did not inhibit infectivity when only added 1 and 2 hours post-infection (hpi). This observation was similar for the entry blocker, hydroxychloroquine, suggesting that the kite-shaped molecules are also entry blockers. The exceptions were chlorprothixene and chlorpromazine, which were still able to reduce infectivity to 45% and 35%, respectively, when added at 1hpi. However, these reductions in infectivity were still much lower than the complete inhibition of infectivity observed for the reverse transcriptase inhibitor, tenofovir. Altogether, these results suggest that the kite-shaped molecules inhibit a SARS-CoV-2-specific entry step.

**Figure 7:**
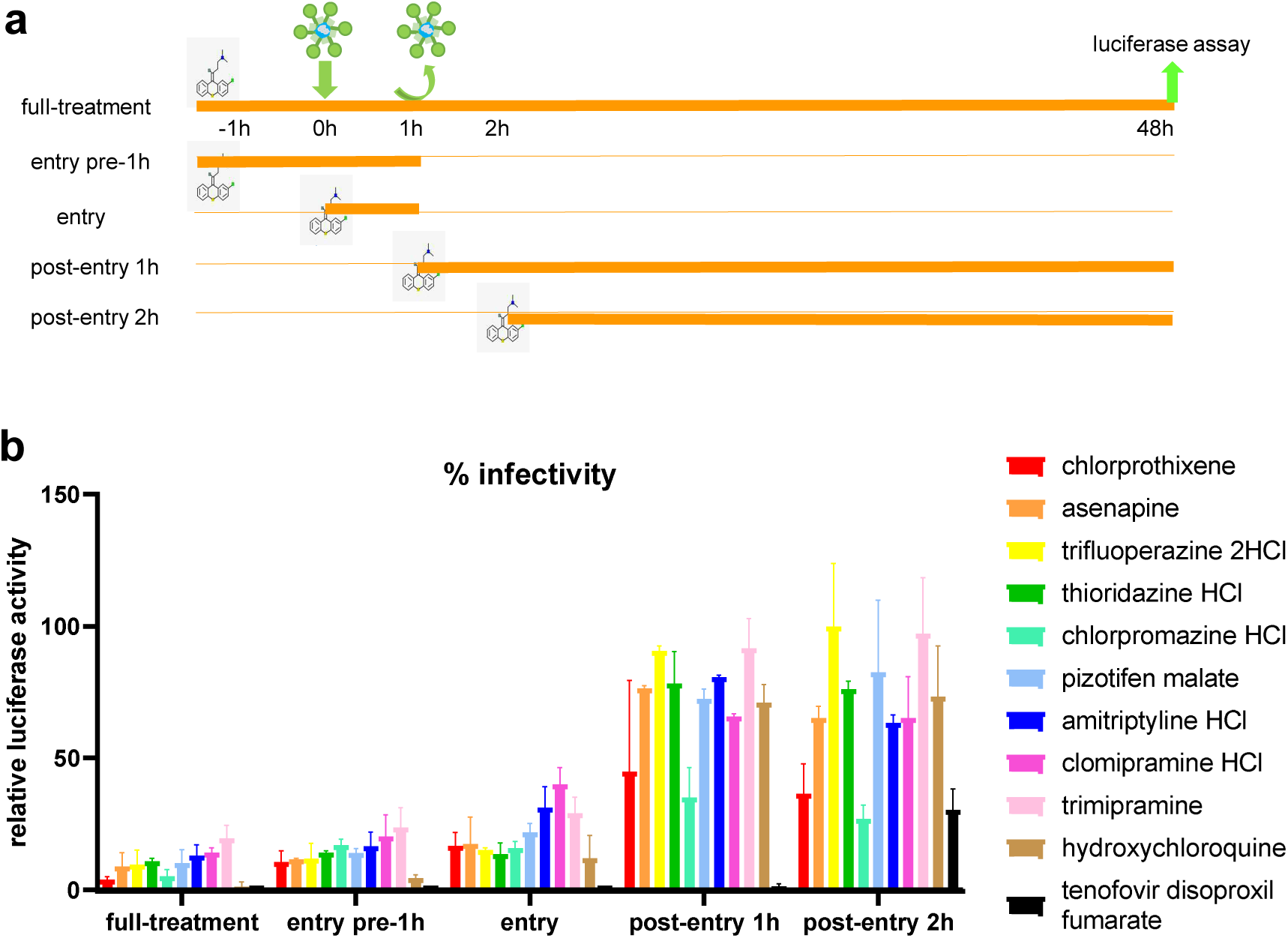
Kite-shaped molecules inhibit SARS-CoV-2 pseudovirus at entry steps. Mouse leukaemia virus pseudotyped with spike protein (S) from severe acute respiratory syndrome coronavirus-2 was used to infect 293T-ACE2 cells in 96-well plates for 48h in a time-of-addition experiment. (a) Schematic of time-of-addition experiment. Full-time treatment involved 1h drug pre-treatment and 1h infection in the presence of drug followed by drug and virus wash-off and addition of fresh drug for the rest of 48h. Entry assay involved 1h infection in the presence of drug with and without 1h drug pre-treatment. The drug and virus were washed off and fresh medium was added without drug for the rest of 48h. Post-entry assay involved no drug pre-treatment and infection in the absence of drug. Following virus wash-off, drug was added at 1 hour post-infection (hpi) or 2hpi for the rest of 48h. (b) Infectivity was measured as luciferase activity and expressed as % infectivity to infected, own solvent control (dimethylsulphoxide, ethanol or water) at the same time-point. Data are presented as mean +/− SD of two repeats.

### Kite-shaped molecules inhibit a post-attachment step

To further delineate the entry step that is inhibited by the kite-shaped molecules, we undertook temperature shift experiments to distinguish between attachment and post-attachment steps that were assumed to proceed (virus attachment) - or not proceed (virus entry, post attachment) - at the low temperature (4°C) employed in the first experiment (Fig. 8a). With this assay, hydroxychloroquine did not inhibit attachment, in agreement with its main role as a post-attachment entry blocker (Fig. 8b). Most of the kite-shaped molecules reduced infectivity to 48-71% at the attachment step apart from chlorprothixene which did not inhibit virus attachment. Thioridazine reduced infectivity to 28% in the attachment assay. These data suggest that all the kite-shaped molecules may reduce attachment to some extent, although it should be acknowledged that this interpretation of the data depends on complete removal of the added drugs at the wash step, which may be dependent on water solubility at 4°C (Supplementary Table S2). Overall, the data suggest that the kite-shaped molecules only modestly inhibit attachment of virus to target cells. In contrast, the kite-shaped molecules significantly reduced infectivity under conditions permissive for the post-attachment, entry step (Fig. 8b), and to levels similar to that of the post-attachment entry blocker control, hydroxychloroquine, suggesting that the kite-shaped molecules are mainly targeting post-attachment entry. These experiments suggest that the kite-shaped molecules are mainly post-attachment entry blockers although some attachment blocking activity cannot be ruled out completely with the current assays employed.

**Figure 8:**
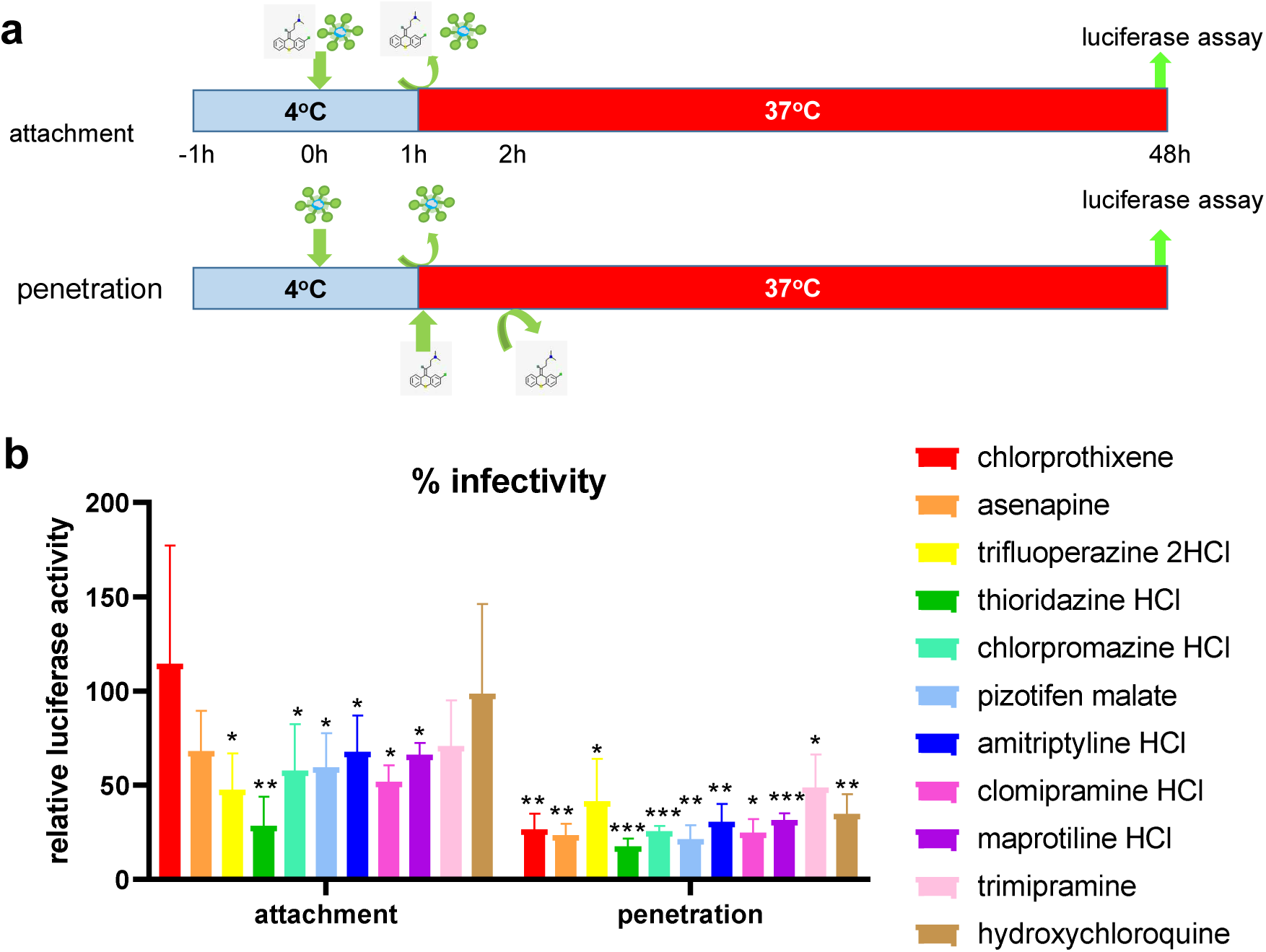
Kite-shaped molecules inhibit SARS-CoV-2 pseudovirus at post-attachment steps. Mouse leukaemia virus pseudotyped with spike protein (S) from severe acute respiratory syndrome coronavirus-2 was used to infect 293T-ACE2 cells in 96-well plates for 48h in a temperature shift experiment. (a) Schematic of temperature shift experiment. Cells were pre-cooled for an hour. In the attachment assay, 1OμM drug diluted in pre-cooled virus aliquots were added to infect for an hour on ice. After 1h, drug and virus were washed off and cells rinsed 3x with cold medium. Fresh, warm medium was added and cells incubated for the remaining 48h at 37°C. In the penetration assay, pre-cooled virus without drug was added to cells at 4°C. The virus was washed off after 1h and cells rinsed 3x with ice-cold medium. Drug in warm medium was then added to incubate with cells for 1h at 37°C. The drug was then washed off with warm PBS (without Mg^2+^ and Ca^2+^) and rinsed with warm medium. Fresh, warm medium was added to continue incubation for the rest of 48h. (b) Infectivity was measured as luciferase activity and expressed as % infectivity to infected, own solvent control (dimethylsulphoxide, ethanol or water). Data are presented as mean +/− SD of four repeats for attachment assays and three repeats for penetration assays. Statistically significant differences are represented by * p<0.05, **p<0.01 and ***p<0.001.

### Asenapine and hydroxychloroquine show additive, but not synergistic effects on SARS-CoV-2 pseudovirus infection

We tested whether asenapine (IC_50_ 1.4μM) and hydroxychloroquine (IC_50_ 0.7μM) had synergistic actions in inhibiting SARS-CoV-2 infectivity by measuring the dose-response behaviour in the assay using a matrix of concentrations of the two drugs. There was no clear indication of any synergistic effects; rather the data implied that the two drugs had additive effects on viral infectivity at low concentrations (Supplementary Information Fig.S2). These data are consistent with the ideas discussed above, that the kite-shaped drugs and hydroxychloroquine both act at the entry steps of the pseudovirus.

## DISCUSSION

Using an MLV backbone pseudotyped with SARS-CoV-2-S we have successfully identified a class of kite-shaped molecules of TCAs with anti-viral activity and IC_50_ in the μM range. Chlorprothixene, one of the top hits in this study, was identified in a repurposing study screening 8,810 drugs that were either FDA-approved or investigational (36).

Methotrimeprazine and piperacetazine also emerged as top hits from that study, and these two compounds share the basic kite-shaped structure of the TCAs. Evidence from observations of patient populations has suggested there was a lower incidence of symptomatic and severe SARS-CoV-2 problems in psychiatric patients (37), and this report was followed up with an *in-vitro* demonstration of the anti-SARS-CoV-2 activity of chlorpromazine (38). Our top 14^th^ hit, trimipramine, has also been identified to cross-inhibit native SARS-CoV-2 infection of Vero E6 in an anti-viral screen using SARS-CoV-S pseudotyped viruses (26). Our top 5^th^ hit, chlorpromazine, has been shown to inhibit infectivity of SARS-CoV-2 in Vero E6 and A549-ACE2 cells and has entered into a clinical trial in France (37,38). Some of our top hits, chlorprothixene, asenapine, thioridazine, amitriptyline, maprotiline, imipramine have been shown to inhibit native SARS-CoV-2 infection, altogether showing the robustness of our pseudotyped system in quantitative, anti-viral drug screening (36) (39) (40) (41).

Many of the kite-shaped molecules selected are TCAs, bind to brain-located receptors and are currently used to treat neurological problems (42). The IC_50_ values reported in Figure 5 may be considered modest by modern criteria (43); for example, peak serum concentrations (Cmax) of the selected drugs within current drug treatment regimes as listed in PubChem database are in the region of 5nM to 1-2μM, with the highest Cmax values being for chlorprothixene (1.4μM). Large variability in Cmax may also arise from differences in drug metabolism and clearance within patient populations (44). For comparison, hydroxychloroquine, which is employed as an anti-malarial and in immunosuppression reaches Cmax values of around 0.4μM but has so far failed to fulfil early promise as a SARS-CoV-2 antiviral (45).

TCAs are known to bind to their neurotransmitter receptors in deep binding pockets within their transmembrane domains composed of 7 transmembrane helices, as exemplified in the 3D structures of drug/receptor complexes (4M48 - nortriptyline/D2 dopamine receptor (46); 3RZE - doxepin/H1 histamine receptor (47)). For the serotonin transporter, a similar binding site exists for citalopram (5I74) (48), a drug that lacks the central cyclic ring of the TCAs, but is otherwise very similar in its 3D structure to the kite-shaped molecules in its binding mode. A different binding site exists at an allosteric site in the pentameric Cys-loop receptor which can bind chlorpromazine (5LG3) (49). Similarly, the binding of amitriptyline to poly (ADP-ribose) polymerase-1 (PARP1) displays an entirely different binding site (50). Low affinity binding of clomipramine, thioridazine and imipramine to the Ebola virus glycoprotein has also been reported, and these compounds were also shown to reduce infectivity of a pseudotyped virus system with IC_50_ values in the 8-13μM range (51). The binding site for these compounds does not have an equivalent in the SARS-CoV-2 spike protein, however Ebola virus and SARS-CoV-2 may share a similar entry route into the cell (52). Hence, none of these structural studies provided clear clues as to the likely protein target of the kite-shaped drugs for inhibition of SARS-CoV-2 infectivity, but they do demonstrate that they can bind to a variety of targets.

Perhaps of greater significance is that TCAs and similar drugs can bind and inhibit the SLC6a19 amino acid transporter (53) that is highly expressed in the intestines and kidneys (54). The structure of LeuT, a bacterial homolog of SLC6a19 and other transporters in the SLC6a grouping has been studied in the presence of diverse tricyclic and similar antidepressants including clomipramine and nortriptyline (PDBIDs 4MMA, 4M48), revealing the nature of inhibition and the binding site (55). SLC6a19 is known to form a stable complex with the ACE2 receptor and the SARS-CoV-2 RBD (PDBID 6M17, see also 6M18, 6M1D) (56), and residues involved in binding drugs in LeuT are conserved in the human SLC6a19 protein. Docking of clomipramine, amitriptyline, and the pharmacophore model shown in Figure 6, into the SLC6a19 atomic model was possible (Supplementary Information Figure S3) and this highlighted aromatic residues and H-bond acceptors that may be involved in the binding of the inhibitory TCAs.

Hence one could hypothesise that the kite-shaped molecules are affecting SARS-CoV-2 infectivity in ACE2-overexpressing kidney cells at an early stage in the viral lifecycle, possibly by modifying the behaviour of the RBD/ACE2/SLC6a19 complex. Nevertheless, there are conceptual problems for any SLC6a19-based strategy aimed at reducing SARS-CoV-2 infectivity in the lungs: ACE2 is highly expressed in airway cells but SLC6a19 is not. Airway-expressed homologs of SLC6a19, such as SLC6a14, SLC6a15 and SLC6a20 may be considered as possible replacements for SLC6a19 in the lungs. However for these proteins there is currently no evidence for any direct interaction with ACE2. One of the candidates, SLC6a15, has been reported to interact directly with several SARS-Cov-2 proteins including the M membrane glycoprotein (57,58), and like ACE2, it appears to carry a C-terminal PDZ-binding motif which therefore offers a potential route for an interaction bridged by an unknown PDZ protein. If SLC6a19 is being replaced by a homolog in the airways, then it could be argued that the most likely candidate is SLC6a15.

The above hypothesis is supported by our time-of-addition and temperature shift experiments, which have identified post-attachment as the main target with some inhibition of attachment step. Moreover, the ability of the kite-shaped molecules to inhibit SARS-CoV-S, SARS-CoV-2-S and MERS-CoV-S pseudoviruses with a preferential inhibition for the two SARS-CoV-S pseudoviruses, suggests that they target a pathway shared by the three viruses but may, in addition, discriminate between different receptor-mediated pathways for viral entry.

Although the kite-shaped molecules generally share a common mode of action, they may also possess unique mechanisms of inhibition. Whereas most of the kite-shaped molecules displayed a similar inhibitory pattern of infectivity, chlorprothixene and chlorpromazine showed some discrepancies. Both inhibited the VSV-G pseudovirus to a greater extent than the other drug hits. Both showed some degree of inhibition when added at 1hpi and 2hpi. Whereas all the other kite-shaped molecules displayed some degree of inhibition of attachment, chlorprothixene did not inhibit attachment. Although some of these discrepancies could be accounted for by the relative toxicity of these two drugs, off-target effects and water insolubility of chlorprothixene, we cannot exclude the possibility that they are results of possession of unique targets. Chlorpromazine has been known to inhibit clathrin-mediated endocytosis, which is utilized by the VSV to enter cells, so it is not surprising that it will inhibit VSV-G pseudovirus infection to some extent (59) (60). Imipramine, a parent compound of trimipramine, blocks macropinocytosis-a potential route of viral endocytosis although activity has not been demonstrated in SARS-CoV-2 infection (61). In an anti-viral screen using native SARS-CoV-2, chlorpromazine was added 2h before infection and after infection but was absent during infection, suggesting that chlorpromazine may inhibit SARS-CoV-2 infection by targeting host pathways, placing it in a class of host-targeting agent (38). Three of our top hits, amitriptyline, maprotiline and imipramine have been shown to prevent SARS-CoV-2 infection by inhibiting acid sphingomyelinase, placing them in a class of host-targeting agents (40). We currently do not have enough evidence to propose whether our kite-shaped drug hits are direct-acting antivirals and/or host-targeting agents. Further work will be required to identify the common and unique modes of action of our drug hits in order to facilitate the formulation of a drug cocktail.

Although it is well recognized that SARS-CoV-2 infects lungs, gut and eyes, increasing evidence suggest liver and kidney tropism with kidney predicted to be the most susceptible (62). Hence, the kidney cell line we used in this study is relevant to SARS-CoV-2 infection biology. 293T cells are devoid of TMPRSS2, hence unable to trigger plasma membrane fusion (19). The TMPRSS2 status of A549 cells is unclear with expression detected in some studies but not others (63) (64). However, the A549-ACE2 cells we used in this study are devoid of TMPRSS2 (65). As a result, our anti-viral screening is limited to drug hits that inhibit the endosomal entry pathway. Most viruses employ either the plasma membrane fusion or the endosomal fusion pathway to enter cells (10). SARS-CoV-2 is peculiar in that it can employ either pathway (19). SARS-CoV-2 enters cells by fusion at the plasma membrane when the membrane protease, TMPRSS2, is available. In the absence of TMPRSS2, SARS-CoV-2 has the flexibility to switch to endosomal entry pathway. Endosomal fusion is the major entry pathway for SARS-CoV-2 in ACE2-deficient cells (20). It is, therefore, of paramount importance for an anti-viral regime to be able to target both pathways. Selective targeting of the default membrane fusion pathway may drive the evolution of SARS-CoV-2 into the embrace of the endosomal fusion pathway.

In conclusion, our study has generated a class of kite-shaped molecules that target a potentially conserved post-attachment step of SARS-CoV-2 cell entry which could inform clinical trials in the current crisis. We have also created a pharmocophore that will allow for improvement in drug design as a broad-spectrum antiviral for future pandemics. There have been four influenza pandemics in 100 years (66). The occurrence of pandemics/outbreaks has clustered in the last ~20 years, including the 1997 Hong Kong bird flu scare, 2003 SARS outbreak, 2009 swine flu, 2014 Ebola outbreak, the 2016 Zika global health concern and now the COVID-19 pandemic (67) (68) (69) (70) (66) (1). In 17 years we have had three deadly coronavirus outbreaks: SARS-CoV (2002–2003), MERS-CoV (2012-2013; ongoing sporadic) and SARS-CoV-2 (2019 to date) (71). Reports of new coronaviruses jumping into humans are emerging (72). Predictions of SARS-like outbreaks every 5-10 years seem reasonable and drugs are needed to stop the current economic chaos being repeated. Currently we are responding to, rather than preparing for a pandemic. We need a first line anti-viral defence to reduce the impact of new coronavirus variants while a vaccine is developed. This drug, or class of drugs, would target conserved steps in the viral life cycle and be broad-spectrum and generally applicable to COVID variants of the current and next pandemics.

## EXPERIMENTAL PROCEDURES

### Cells

293T, 293T-ACE2, A549-ACE2, Caco2, Huh-7 and Vero cells were cultured in Dulbecco’s modified Eagle’s medium with 4mM glutamate (DMEM; Sigma) and supplemented with 10% fetal calf serum (FCS; Sigma), 100 units/ml penicillin and 100μg/ml streptomycin (Sigma) at 37°C, 5% CO_2_. The culture medium of A549-ACE2 was supplemented with 1μg/ml puromycin (Sigma). The culture medium of Caco2 and Huh-7 was supplemented with 1x non-essential amino acid. Calu3 cells were cultured in Minimal Essential medium supplemented with 2mM glutamate.

### Pseudovirus system

293T cells seeded at 4×10^6^ per 100mm dish were co-transfected with 6μg of a plasmid encoding MLV gag-pol, 8μg of the transfer vector encoding a luciferase reporter and 6μg of a plasmid encoding an empty vector, a viral envelope glycoprotein SARS-CoV-S, SARS-CoV-2-S, MERS-CoV-S or VSV-G using calcium phosphate (125mM CaCl_2_, 0.7mM Na_2_HPO_4_, 70mM sodium chloride, 25mM Hepes pH 7.05) (see Fig. 1). The medium was replaced with fresh medium after 24h and supernatant containing pseudoviruses was harvested after 48h, clarified by centrifugation at 1000rpm/4°C for 10min, filtered through 0.45μm filter and stored at −80°C. The resulting pseudovirus contains an MLV gag-pol backbone packaging a luciferase reporter genome and displaying one of the viral envelope glycoproteins. The MLV pseudotyped with an empty vector is bald.

### Anti-viral drug screening

Two libraries from APExBIO containing 1363 FDA-approved drugs (cat:L1021) and 137 natural compounds (cat:L1039) were used in screening. Drug stocks at 10μM were diluted into 1μM in their own solvents or diluted directly into medium to 20μM. 293T-ACE2 cells seeded at 25,000 cells per well of 96-well plates were pre-treated with 10μM of individual drugs, in duplicate, for 1h. After 1h, 25μl of pseudovirus was added together with drugs to maintain the final drug concentration at 10μM. A parallel set of 96-well plates were set up with only drugs without pseudovirus, in duplicate, to test for drug cytotoxicity. After incubation for 37°C, 5%CO_2_ for 48h, they were tested for luciferase activity for % infectivity relative to the infected, solvent controls and for % viability relative to the solvent controls. For the generation of concentration curves, serial dilutions of drugs were titred, in duplicates and IC_50_ was calculated using Prism9 (GraphPad).

### Luciferase assay

Cells were lysed by the addition of 100μl of passive lysis buffer (Promega) to each well and shaken for >15min. Luciferase assay was carried out as described in (73,74) (75) in a buffer containing 0.0165 M glycylglycine, 0.01 M MgSO_4_, 2.65 mM EGTA, 10.5 mM potassium phosphate, 1.4 mM adenosine 5’-triphosphate, 0.86 mM dithiothreitol (DTT), 0.175 mg/ml bovine serum albumin, and 0.035 mM luciferin (Promega) using 50μl of the lysate and a luminometer (Berthold Technologies, Germany).

### XTT viability assay

Cell viability was measured by addition of 50μl of 1mg/ml 2,3-Bis-(2-Methoxy-4-nitro-5-sulfophenyl)-2H-tetrazolium-5-carboxanilide, disodium salt (XTT, Biotium) and 20μM N-methyl dibenzopyrazine methyl sulfate (Cayman) in culture medium to each well for 2-4h at 37°C/5% CO_2_. Absorbance was read at 450nm with a reference wavelength of 650nm using a plate reader (Bio-Tek Synergy HT).

### Time-of-addition experiment

In time-of-addition experiment, drugs were added at different times of infection (see Fig. 7a). 25μl of pseudovirus were added to each well for 1h, then washed off with 2x PBS. Inhibition of the entry steps was assayed by the addition of drugs during the 1h infection with and without an 1h pre-treatment. Drugs were washed off together with the virus at 1hpi and incubation continued until 48hpi in the absence of drugs. Inhibition of post-entry steps involved no drug pre-treatment and infection in the absence of drug. After washing off the virus at 1hpi, drugs were added immediately or at 2hpi and were present for the duration of the rest of the 48h infection. A full-time treatment was included as a control in which drugs were added 1h pre-infection and during infection. After washing off the drugs and viruses at 1hpi, fresh drugs were added until 48hpi.

### Temperature shift experiment

Cells seeded on 96-well plates were pre-cooled on ice for 1h (see Fig. 8a). For the attachment assay, drugs were diluted into pre-cooled viruses on ice to 10μM just before infection and the 100μl virus-drug mix was added to each well for 1h. After 1h, the wells were washed 3x with ice-cold medium. Warm medium was added and incubation continued at 37°C/5%CO2 until 48hpi. For the penetration (post-attachment) assay, 100μl of pre-cooled virus without drugs were added to each well for 1h infection. After 1h, the wells were washed 3x with ice-cold medium. Drugs diluted in warm medium were added and incubate at 37°C, 5%CO_2_ for 1h. After 1h, the drugs were washed off with 1x warm PBS (without Mg^2+^, Ca^2+^) and 1x with warm medium. Fresh, warm medium was added and incubation continue at 37°C, 5%CO_2_ until 48hpi.

### Western blotting

Western blotting was performed as described (73,74,76–78). Protein lysates were harvested into radioimmunoprecipitation assay buffer (RIPA) buffer (50mM Tris pH8.0, 150mM NaCl, 1% NP40, 0.5% Na deoxycholate, 0.1% SDS) plus protease (Sigma) and phosphatase inhibitors (APExBIO). Proteins from equal number of cells were separated on TGX Stain-Free SDS-PAGE gel (Bio-Rad), transferred to polyvinylidene difluoride membrnaes (Millipore), blocked in 5% semi-skimmed milk (Marvel) in 0.1% Tween 20 (Sigma)/TBS (50mM Tris pH 7.4, 150mM NaCl) before being probed against primary and horseradish peroxidase (HRP)-conjugated secondary antibodies in blocking buffer. Anti-ACE2 antibody (Proteintech) was used at 1:2000 and anti-mouse HRP (Cell Signaling Technology) at 1:1000. Anti-SARS-CoV-2 spike antibody (BEI Resources NR-52947) was used at 1:1000 and anti-rabbit HRP (Cell Signaling Technology) at 1:1000. Protein bands were detected using Clarity^TM^ ECL substrate (Bio-Rad). Images were captured and quantified using ChemiDoc^TM^ XRS+ system (Bio-Rad) and ImageLab 6.0.1 software (Bio-Rad).

### Pharmacophore & Docking Studies

A pharmacophore was built using the pharmacophore query module implemented in Molecular Operating Environment (MOE) (79). In brief, Compounds within 0.5 log fold activity of the highest inhibitory activity were considered active whereas compounds yielding infectivity ranging above 100% infectivity (control) values were considered inactive. These compounds were stochastically searched for conformers using default parameters and packed as a known dataset during pharmacophore development. Docked conformations of asenapine due to its fused ring rigid structure were considered as templates for pharmacophore development. The pharmacophoric features were calculated using AutoPH4 script (80) with holo conditions, and manually optimized for maximum performance. The final model was used on the complete drug dataset to access the screening performance.

Docking studies were performed using GOLD software version 2020.3.0 (81). in summary, 2D depicted structures of compounds were extracted from the PubChem database (82) and were compiled in a MOE database. The dataset was washed for adjuvant atoms and protonated at pH 7.4. Partial charges were computed using MMFF94x methods as implemented in MOE (79). The charge calculated ligands were energy minimized for relaxed three-dimensional conformations. The LeuBAT structure with clomipramine (4MMD) (83) and the SLC6a19 structure (6M18) (84) were downloaded from the PDB database. The structural discrepancies in the models were corrected using the structure correction module implemented in MOE. The corrected structures were protonated, energy minimized and saved for docking simulations. Both protein and ligands were considered flexible during the simulations. To explore diversity of conformational solutions, a 15Å area around the ligand binding site was selected. Furthermore, the cavity was strictly restricted to solvent accessible area using LIGSITE implemented in GOLD. To ensure reproducibility a total of 100 conformations were generated and ranked according to the scoring function.

### Statistical analysis

Statistical analysis was performed and graphs were plotted using Prism 9.0 (GraphPad). Shapiro-Wilk normality test and one sample t-test were used for the analysis of temperature shift data against a theoretical mean of 100. A p value of <0.05 was considered statistically significant.

Data availability: The pharmacophore and docking results may be obtained from the corresponding author. The authors declare that they have no conflicts of interest with the contents of this article

*This article contains Supporting Information*.

