## Supplemental information for "Kite-shaped molecules block SARS-CoV-2 cell entry at a post-attachment step"

**Running title: Repurposing for coronaviruses.**

**Shiu-Wan Chan*, Talha Shafi and Robert C. Ford**

**Faculty of Biology, Medicine and Health, School of Biological Sciences, The University of Manchester, Michael Smith Building, Oxford Road, Manchester M13 9PT, United Kingdom**

**Supplementary information**

**Table S1**

**Table S2**

**Figure S1**

**Figure S2**

**Figure S3**

| Table S1 Kite-shaped molecules in order of inhibition of pseudovirus infectivity | | | | |
|---|---|---|---|---|
| | % infectivity[a] | % viability[b] | 2D structure | 3D structure |
| chlorprothixene | 4.1 | 97 | 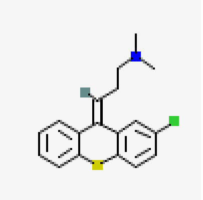 | 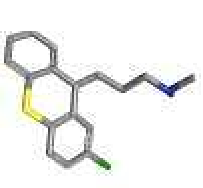 |
| asenapine | 4.38 | 92 | 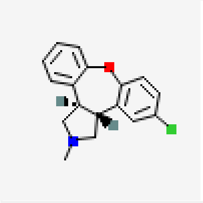 | 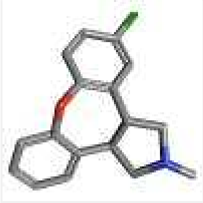 |
| Trifluoperazine 2HCl | 4.84 | 83 | 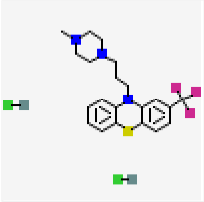 | 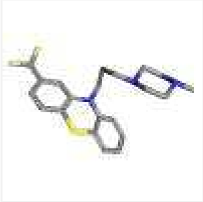 |
| thioridazine HCL | 5.98 | 71 | 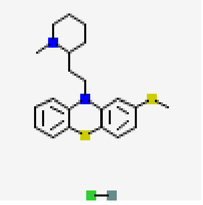 | 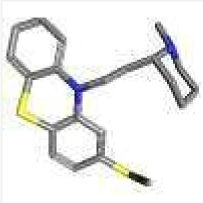 |
| Chlorpromazine HCl | 6.24 | 65 | 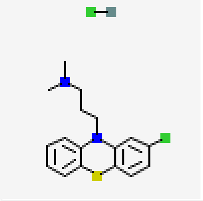 | 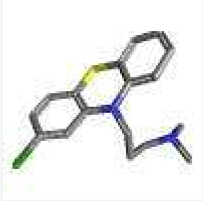 |
| Pizotifen Malate | 7.01 | 106 | 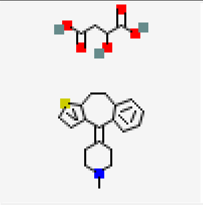 | 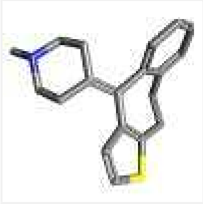 |

| | | | | |
|---|---|---|---|---|
| Amitriptyline HCl | 9.05 | 112 | 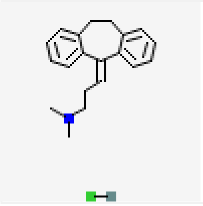 | 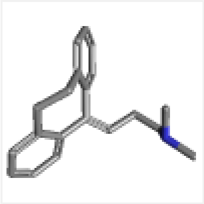 |
| Clomipramine HCl | 9.23 | 115 | 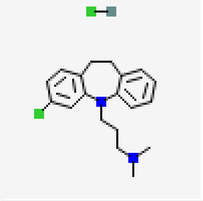 | 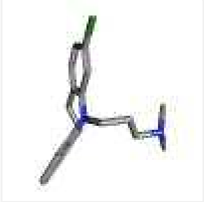 |
| maprotiline hcl | 9.38 | 99 | 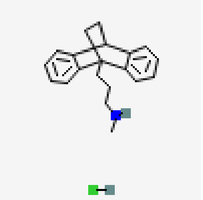 | 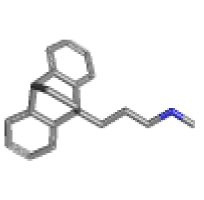 |
| cyclobenzaprine HCl | 9.49 | 101 | 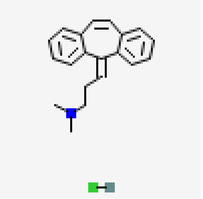 | 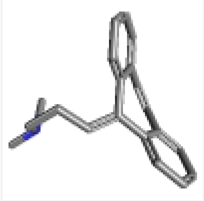 |
| Desloratadine | 10.25 | 99 | 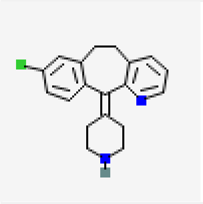 | 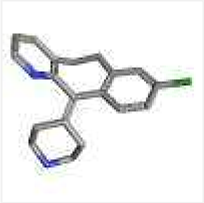 |
| promethazine HCl | 11.32 | 99 | 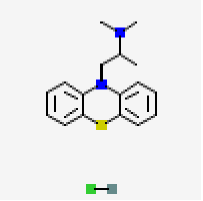 | 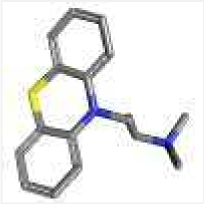 |
| Carvedilol | 13.1 | 81 | 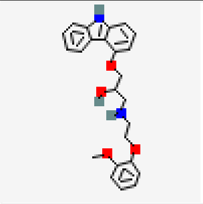 | 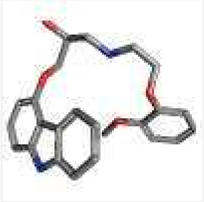 |

| | | | | |
|---|---|---|---|---|
| trimipramine | 13.54 | 92 | 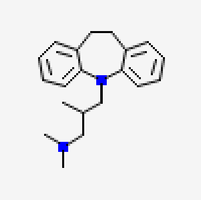 | 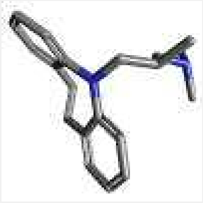 |
| MLN8237 | 15.36 | 63 | 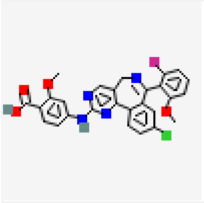 | 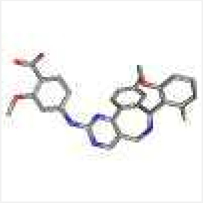 |
| S/GSK1349572 | 16.2 | 94 | 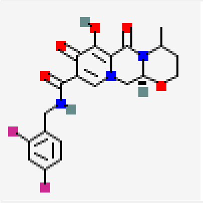 | 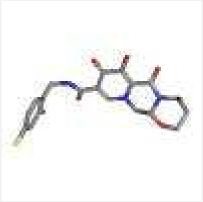 |
| Imipramine- | 16.34 | 99 | 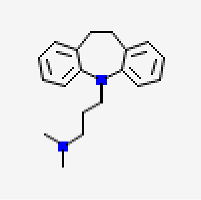 | 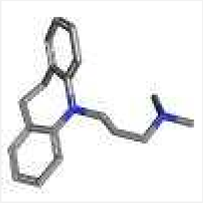 |
| Prochlorperazine | 16.41 | 94 | 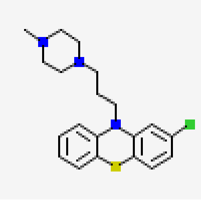 | 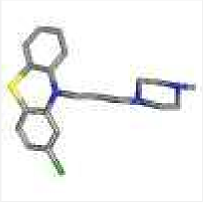 |
| Cyproheptadine hydrochloride | 16.9 | 94 | 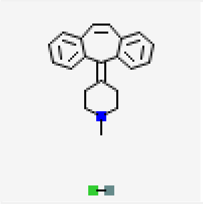 | 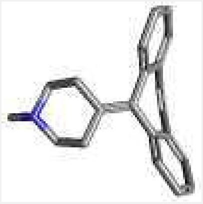 |
| amoxapine | 19.9 | 103 | 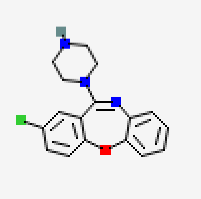 | 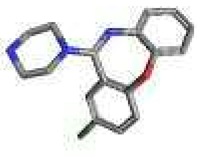 |

| | | | | |
|---|---|---|---|---|
| Olanzapine | 26.22 | 106 | 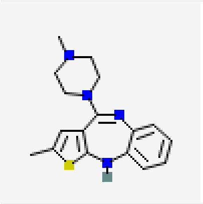 | 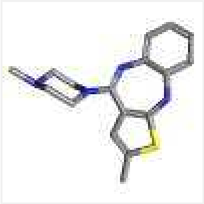 |
| elacridar | 27.04 | 97 | 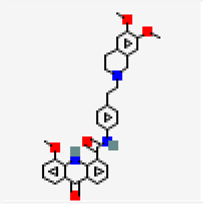 | 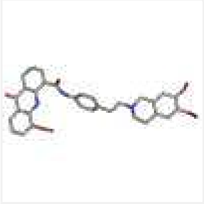 |
| Clozapine | 28.31 | 103 | 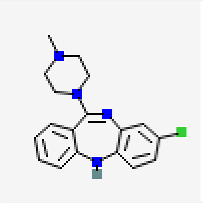 | 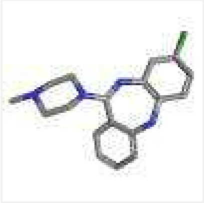 |
| Loxapine Succinate | 35.01 | 94 | 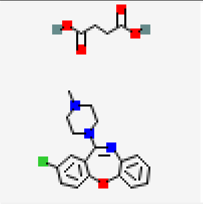 | 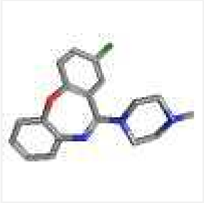 |
| Dehydrocostus Lactone | 36.23 | 90 | 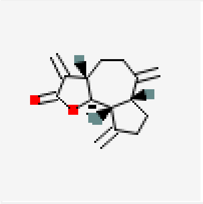 | 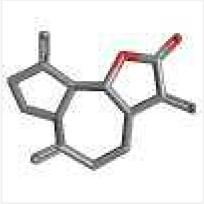 |
| Rupatadine Fumarate | 38.01 | 74 | 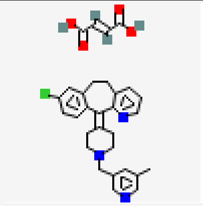 | 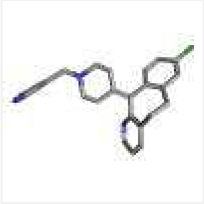 |
| mianserin Hcl | 41.95 | 100 | 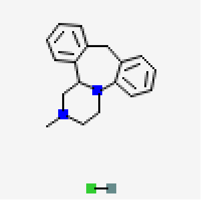 | 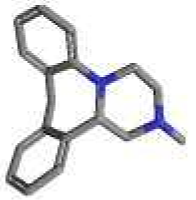 |

| | | | | |
|---|---|---|---|---|
| Clofazimine | 45.61 | 77 | 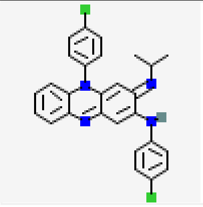 | 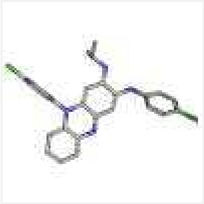 |
| Mesoridazine besylate | 48.73 | 97 | 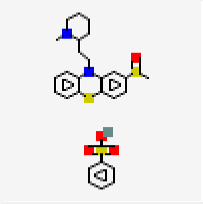 | 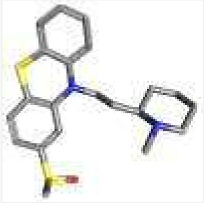 |
| conivaptan hcl | 51.97 | 102 | 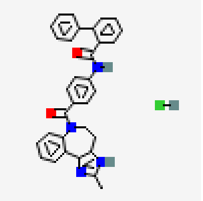 | 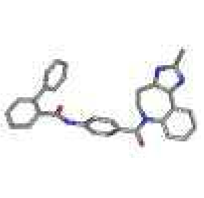 |
| Tacrine hydrochloride | 57.04 | 97 | 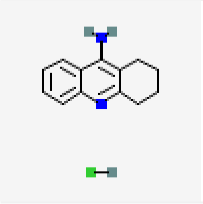 | 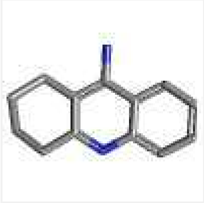 |
| Ketotifen Fumarate | 60.4 | 97 | 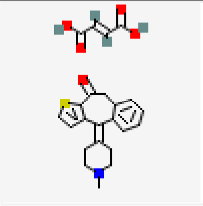 | 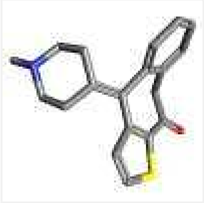 |
| lonafarnib | 64.73 | 111 | 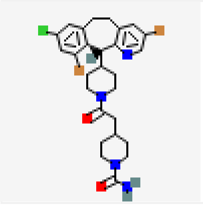 | 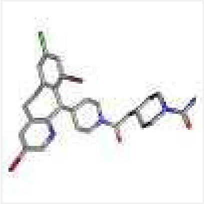 |
| loratadine | 67.15 | 94 | 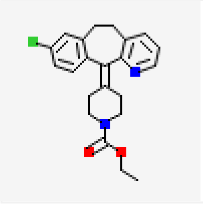 | 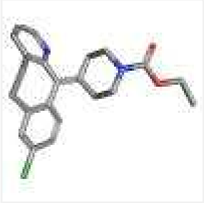 |

| | | | | |
|---|---|---|---|---|
| Diphenhydramine HCl | 67.66 | 101 | 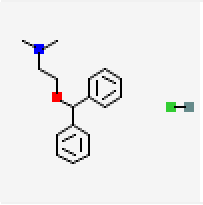 | 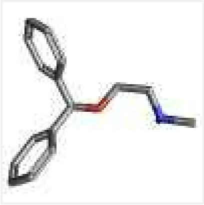 |
| Mirtazapine | 72.63 | 104 | 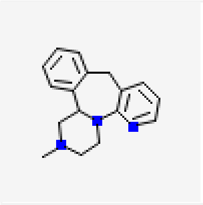 | 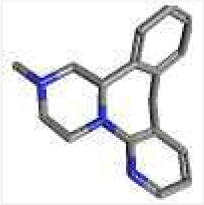 |
| Epinastine HCl | 72.78 | 94 | 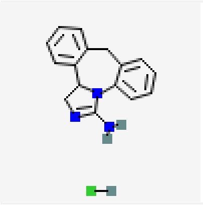 | 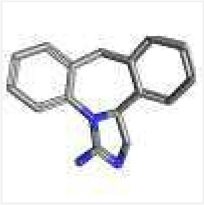 |
| Etodolac | 76.15 | 100 | 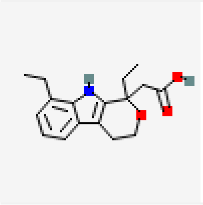 | 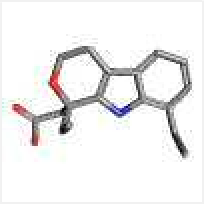 |
| Carprofen | 80.32 | 103 | 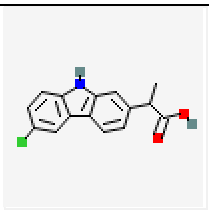 | 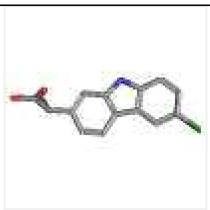 |
| Nevirapine | 80.71 | 103 | 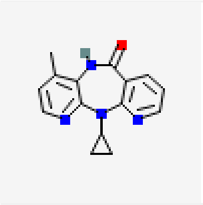 | 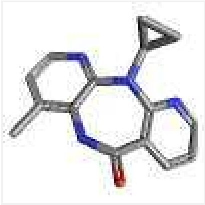 |

| | | | | |
|---|---|---|---|---|
| Carbamazepine | 81.27 | 105 | 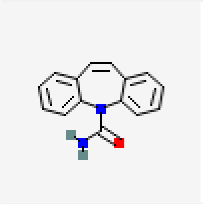 | 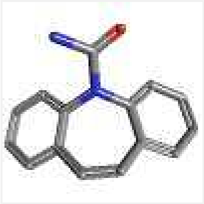 |
| Diacerein | 81.89 | 126 | 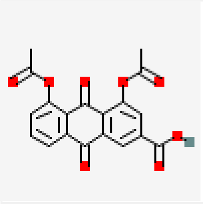 | 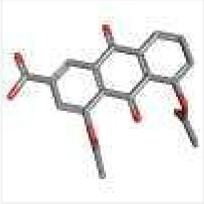 |
| Alcaftadine | 85.44 | 97 | 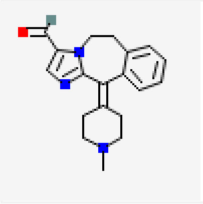 | 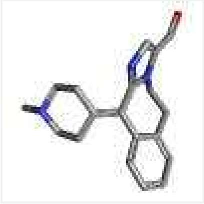 |
| Pranoprofen | 87.46 | 114 | 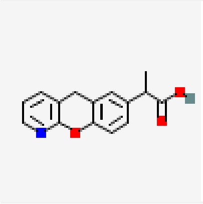 | 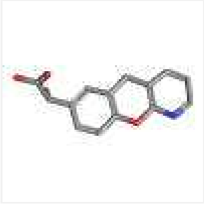 |
| Eslicarbazepine acetate | 90.75 | 102 | 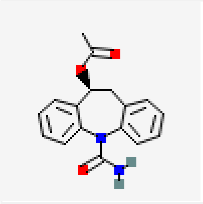 | 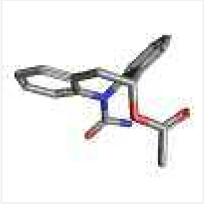 |
| Olopatadine HCl | 91.21 | 99 | 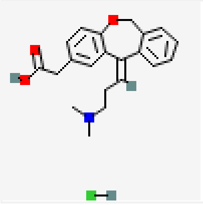 | 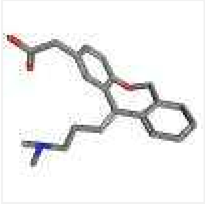 |

| | | | | |
|---|---|---|---|---|
| Dibenzothiophene | 92.11 | 102 | 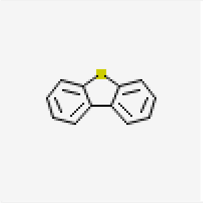 | 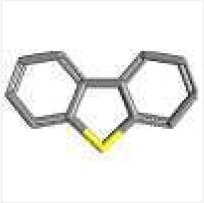 |
| Tianeptine sodium | 94.8 | 102 | 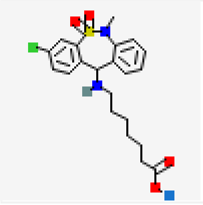 | 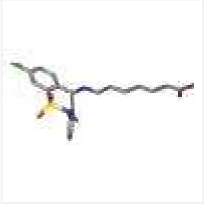 |
| Oxcarbazepine | 95.77 | 109 | 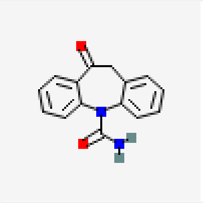 | 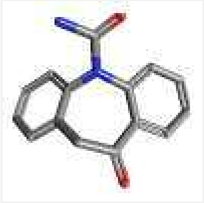 |
| flumazenil | 99.18 | 104 | 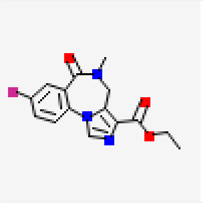 | 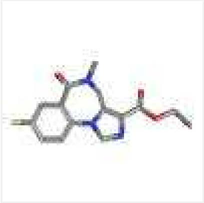 |
| Quetiapine Fumarate | 99.77 | 96 | 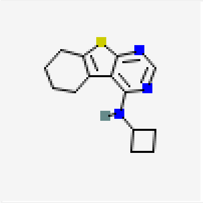 | 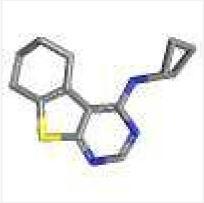 |
| 8-methoxypsoralen | 100.26 | 105 | 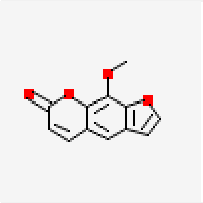 | 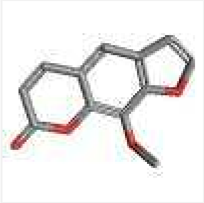 |

| | | | | |
|---|---|---|---|---|
| Ramelteon | 104.18 | 99 | 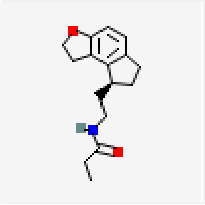 | 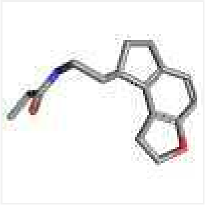 |
| Aloin | 107.91 | 103 | 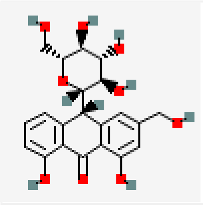 | 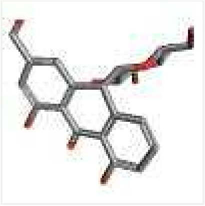 |
| 5-methoxypsoralen | 109.79 | 111 | 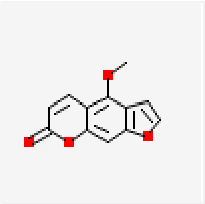 | 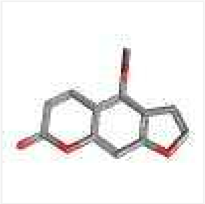 |
| quetiapine | 112.56 | 101 | 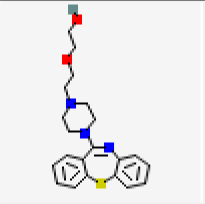 | 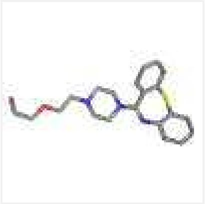 |

a % infectivity relative to the infected, DMSO control

b % viability relative to the un-infected, DMSO control

drug structures obtained from PubChem

Table S2 Water solubility of kite-shaped molecules and hydroxychloroquine

| Drug | water solubility[a] | water solubility[b] |
|---|---|---|
| hydroxychloroquine sulphate | 60 μM | 200mM |
| chlorprothixene HCl | 0.9 μM | 28mM |
| asenapine maleate | 78 μM | 9mM |
| trifluoperazine HCl | 18 μM | 104mM |
| thioridazine HCl | 2.1 μM | 122mM |
| chlorpromazine HCl | 12 μM | 141mM |
| pizotifen malate | 16 μM | 0.3mM[c] |
| amitriptyline HCl | 14 μM | 99mM |
| clomipramine HCl | 41 μM | 71mM |
| maprotiline HCl | 150 μM | 159mM |
| trimipramine maleate | 88 μM | 5mM |

a Values are predictions for the uncharged form obtained from https://go.drugbank.com

b. Values are for the salts and from chemical supplier datasheets, where available.

c. In a 1:8 dimethylformamide:water mixture.

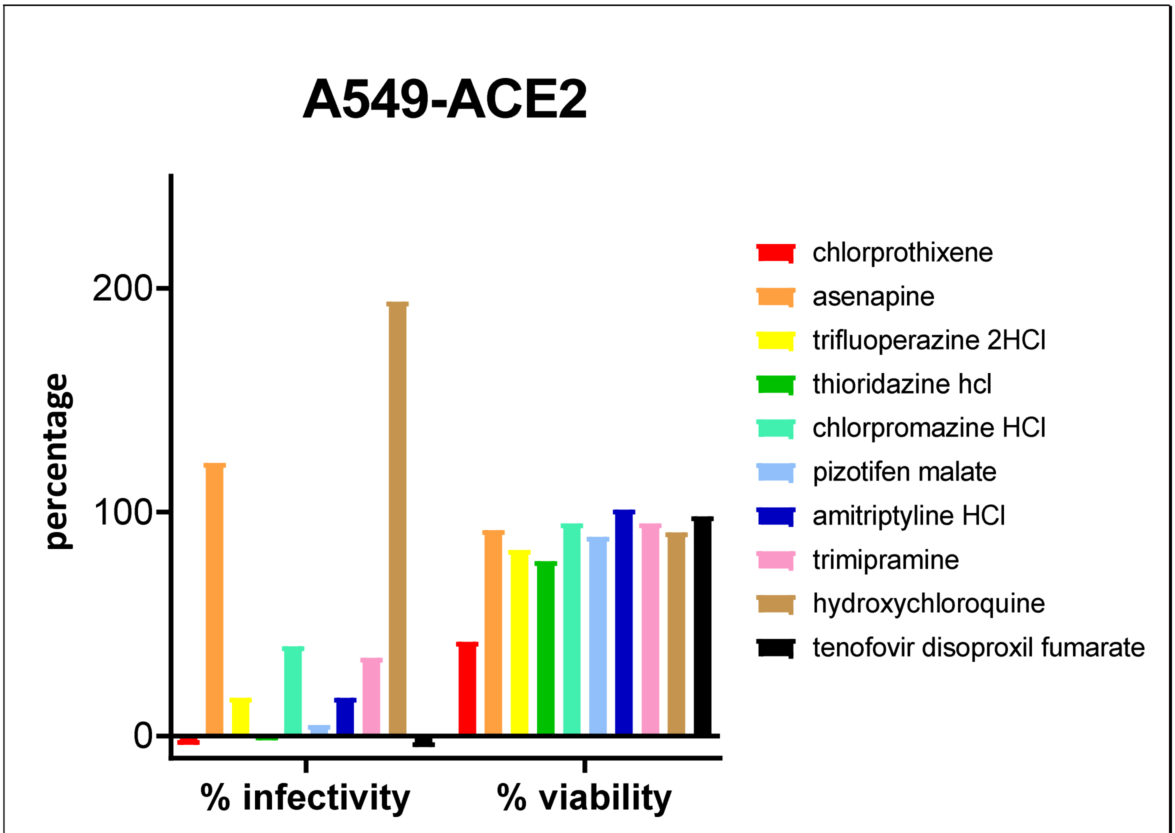


Figure S1

**Fig. S1 The majority of kite-shaped molecules inhibit infectivity of SARS-CoV-2 pseudovirus in A549-ACE2 cells.** Mouse leukaemia virus pseudotyped with spike protein (S) from severe acute respiratory syndrome coronavirus-2 was used to infect 12,000 cells/well A549-ACE2 cells in 96-well plate for 48h. Infectivity was measured as luciferase activity and expressed as % infectivity to infected, own solvent control (dimethylsulphoxide, ethanol or water). Viability was measured by XTT assays in un-infected cells and expressed as % viability to solvent control (dimethylsulphoxide, ethanol or water). Data are from one repeat

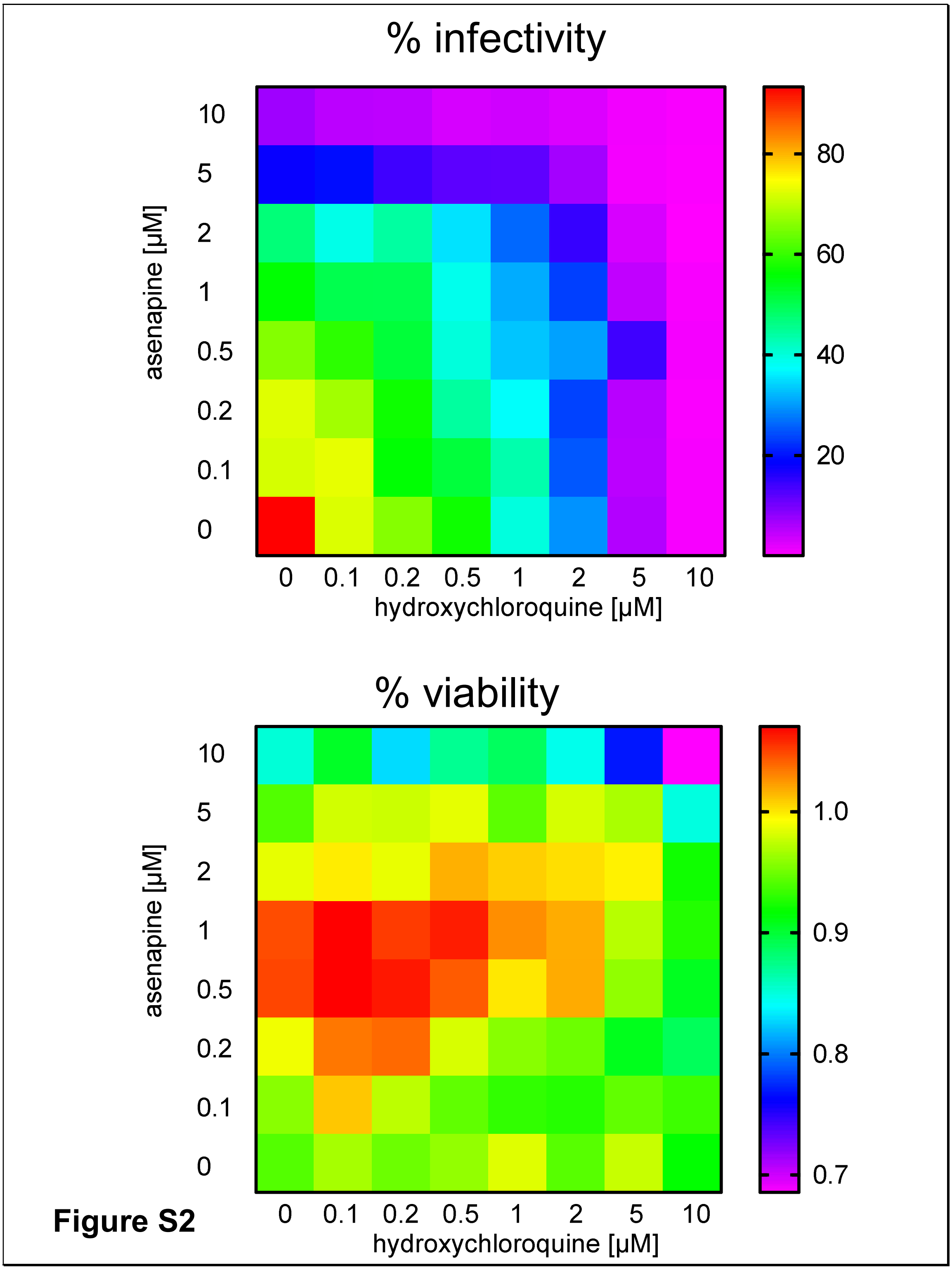


**Fig. S2 Asenapine and hydroxychloroquine show additive inhibitory effect on SARS-CoV-2 infectivity.** Mouse leukaemia virus pseudotyped with spike protein (S) from severe acute respiratory syndrome coronavirus-2 was used to infect 293T-ACE2 cells in 96-well plate for 48h in the presence of serial doses of the asenapine and hydroxychloroquine, as indicated, with 1h pre-treatment. Infectivity was measured as luciferase activity and expressed as % infectivity to infected, own solvent control (dimethylsulphoxide or water). Viability was measured by XTT assays in un-infected cells and expressed as % viability to solvent control (dimethylsulphoxide or water). Date are presented as heat maps generated using Prism9.0 (GraphPad). Data are from one repeat.

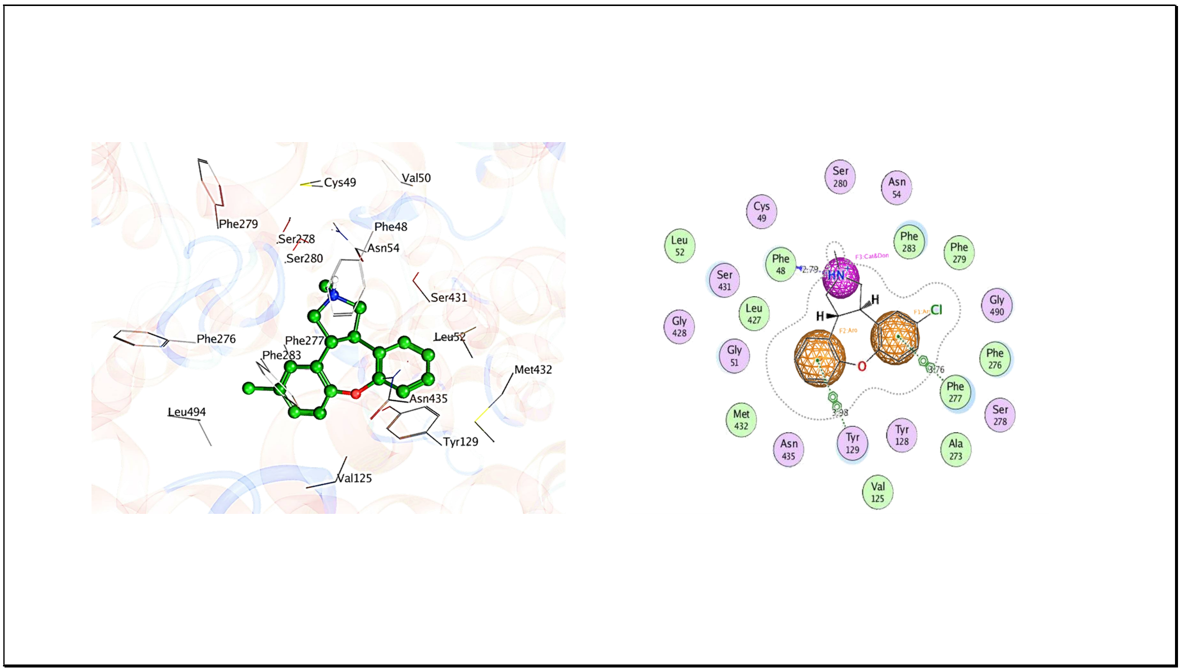


**Supplementary Figure S3: diagrammatic representation of ligand- protein interaction and pharmacophore mapping.** A) Ligand protein profile of the highest ranked conformation of asenapine within the SLC6a19 binding cavity. B) two dimensional representation of ligand protein interaction of asenapine overlaid with the pharmacophore model.
